# Immunisation of chickens with inactivated and/or infectious H9N2 avian influenza virus leads to differential immune B cell repertoire development

**DOI:** 10.1101/2024.07.09.602583

**Authors:** Stefan Dascalu, Joshua E. Sealy, Jean-Remy Sadeyen, Patrik G. Flammer, Steven Fiddaman, Stephen G. Preston, Robert J. Dixon, Michael B. Bonsall, Adrian L. Smith, Munir Iqbal

## Abstract

Avian influenza viruses (AIVs) are a major economic burden to the poultry industry, and pose serious zoonotic risk, with human infections being reported every year. To date, the vaccination of birds remains the most important method for the prevention and control of AIV outbreaks. Most national vaccination strategies against AIV infection use whole-virus inactivated vaccines, which predominantly trigger a systemic antibody-mediated immune response. There are currently no studies that have examined the antibody repertoire of birds that were infected with and/or vaccinated against AIV. To this end, we evaluate the changes in the H9N2-specific IgM and IgY repertoires in chickens subjected to vaccination(s) and/or infectious challenge. We show that a large proportion of the IgM and IgY clones were shared across multiple individuals, and these public clonal responses are dependent on both the immunisation status of the birds and the specific tissue that was examined. Furthermore, analysis reveals specific clonal expansions which are restricted to particular H9N2 immunisation regimes. These results indicate that both the nature and number of immunisations are important drivers of the antibody responses and repertoire profiles in chickens following H9N2 antigenic stimulation. We discuss how the repertoire biology of avian B cell responses may affect the success of AIV vaccination in chickens, in particular the implications of public versus private clonal selection.

## 1. Introduction

Avian influenza viruses (AIVs) are responsible for major economic losses to the poultry sector and a detrimental impact on the health and welfare of chickens (1). AIVs can also transmit between species, causing a disease burden in other livestock, wild animals and humans, emphasising the importance of infection control in poultry (2). Prevention of poultry outbreaks is facilitated using biosecurity practices and vaccination. However, the efficacy of vaccines is frequently hindered by the high mutation rate of influenza genes that results in antigenic drift (3).

Influenza viruses are classified based on their haemagglutinin (HA) and neuraminidase (NA) surface proteins, which mediate cellular attachment and release respectively (2). These proteins also serve as major antigenic targets for the adaptive immune systems of avian and mammalian hosts. Indeed, the majority of the antibody responses are directed towards the HA and NA proteins, with protective titres being able to block the infection of cells (4). The hyperdiverse ”anticipatory” repertoire of BCR/Ig affords the ability of B cells to recognise a wide array of non-self antigens and this is created by a combination somatic rearrangement, junctional modification, pairing of heavy and light chains and, in birds, gene conversion, which occurs in the bursa of Fabricius (5,6).The critical antigen binding region of the rearranged Ig is known as the complementary determining region 3 (CDR3) (7).

High-throughput sequencing (HTS) is increasingly being used to characterise human and mouse antibody repertoires (8,9). Such studies have allowed for the identification of antigen-specific antibody CDR3 rearrangements reactive against dengue virus, influenza virus, human immunodeficiency virus (HIV), tetanus toxoid, *H. influenzae*, group C meningococcal polysaccharides, and SARS-CoV-2 virus (10–13). HTS applied to B cell repertoires also facilitate the detection of B-cell malignancies, understanding autoimmunity, and characterising vaccine responses (8). As such, HTS analysis of the B cell repertoire in response to different HIV vaccination regimes enabled the identification of a vaccine candidate with superior performance compared to alternatives (10).

To date, limited information is available regarding the dynamics of antibody repertoires of birds, especially in the context of infection or vaccination. Poultry, in particular commercial chickens, are subject to an intensive vaccination schedule starting as early as day 1 post-hatch, employing up to 16 or so vaccines over the 5 to 7 week production period of broiler (meat) chickens (14). Many of the AIV vaccines are based on inactivated virus formulations which predominantly stimulate humoral immunity (4). As humoral responses are also known to contribute significantly to protection against AIVs, we set out to explore the effects of vaccination and/or infection on the IgM and IgY repertoires of chickens.

## 2. Materials and Methods

### 2.1. Viruses and vaccines

Recombinant reverse-genetics (RG) viruses were generated using established methods (15). RG viruses comprised all eight genes from the A/chicken/Pakistan/UDL-01/2008 (UDL- 01/08) influenza virus. Influenza virus vaccines were prepared by first inactivating stocks with β-propiolactone 0.1% (v/v) at room temperature for eight hours, then incubating at 4°C for 24 hours (16). Virus inactivation was confirmed through three sequential inoculation experiments of 10 day-old embryonated hen’s eggs. Inactivated virus was then concentrated by ultracentrifugation at 135,000 xg for two hours. Virus was resuspended in PBS and titrated by haemagglutination (HA) assay.

### 2.2. Experimental design and tissue samples

White leghorn chickens (Valo breed, n=70) were hatched and reared in the poultry facility at the Pirbright Institute according to national and institute-specific regulations (licence number P68D44CF4). Birds were divided into 6 treatment groups (G1-6) and maintained together until infection, then transferred to isolators at day 21 post-hatching. Uninfected birds remained together for the duration of the experiment. Chickens from G1 (n=10), G2 (n=10), G4 (n=10), and G5 (n=10) received a single subcutaneous injection of 0.2ml of 1024 HAU/dose inactivated H9N2 vaccine immediately after hatching. Individuals from G2 and G5 received an additional dose of the vaccine at 14 days after hatching. Birds from G3 (n=15) and G6 (n=15) did not receive the vaccine. Weekly blood samples were collected from all birds immediately after the first intervention (i.e. infection or vaccination). At day 21 post-hatching, birds from G1-3 received a single intranasal inoculation of 0.1ml of 10^6^ pfu/ml H9N2 low pathogenic avian influenza virus (50 µl in each nostril). Following infection, birds (G1-3) were weighed daily and buccal and cloacal swabs were collected each day for 10 consecutive days to confirm infection status via qRT-PCR (see below). At day 3 and at day 14 post-infection, 5 birds from each group (G1- G6) were culled and tissue samples (trachea, spleen and bursa) were harvested and stored in RNAlater (Thermo Fisher Scientific). Additionally, at day 7 post-infection, 5 birds from G3 and G6 were culled and their tissues were harvested as for the rest of the time points. A diagram of the experimental design is provided in Supplementary Figure 1. The tissues used for repertoire analysis were the ones harvested at the last time point of the experiment (14 days post-infection).

### 2.3. Haemagglutination and haemagglutination inhibition assays

The HA titre of virus and the haemagglutination inhibition (HI) titre of chicken antisera were determined using established protocols (17). Briefly, to determine HA titre, a two-fold dilution series of virus in PBS was prepared in V-bottom 96-well plates (Thermo Fisher Scientific) and incubated 1:1 with 1% chicken red blood cells (RBCs) in PBS for one hour at 4°C. The HA titre was recorded as the reciprocal of the highest dilution that virus caused complete haemagglutination of RBCs.

To determine the HI titre, a series of 25 μl two-fold dilutions of post-infection chicken polyclonal antisera was incubated with 4 haemagglutinating units (HAU) per 25 µl of virus for one hour at room temperature then incubated 1:1 with chicken 1% RBCs in PBS for one hour at 4°C. The HI titre was recorded as the reciprocal of the highest dilution of chicken antiserum that completely inhibited haemagglutination of RBCs.

### 2.4. ELISA

To assess the levels of H9N2-reactive IgM and IgY in vaccinated and/or challenged birds, enzyme-linked immunosorbent assays (ELISAs) were developed using inactivated virus. Nunc MaxiSorp (Thermo Fisher Scientific) flat-bottom 96-well plates were coated with 50 µl per well of 0.25 µg/ml inactivated virus and incubated overnight at 4°C. The following day, plates were washed 4 times using 0.1% Tween 20 (Sigma-Aldrich) in PBS. Subsequently, the plates were incubated for 30 minutes at room temperature using 100 µl of blocking buffer (0.25% BSA in 0.1% Tween-20 in PBS). Following another wash step, 50 µl of 1:800 serum dilution was added to duplicate wells and incubated for 1h at room temperature. The plates were then washed again and incubated for 30 minutes with 50 µl of either a 1:7000 dilution of goat IgG conjugated with horseradish peroxidase (HRP) having anti-chicken IgY specificity (Biorad), or 1:3000 dilution of goat IgG conjugated with horseradish peroxidase (HRP) having anti-chicken IgM specificity (Invitrogen, cat. PA1-84676). After another wash step, 50 µl of OptEIA TMB Substrate (BD Biosciences) was added to each well of the plate and incubated at room temperature. The reaction was stopped using 50 µl of 2M H_2_SO_4_ after 20 minutes of incubation. The optical density of the wells was then read using a 450/630 nm setting of a ELx808 plate reader (BioTek). The raw optical density (OD) values were then standardised using the sample to positive ratio. For this, a sample of pooled sera from three 24-day-old naïve birds from a separate experiment was used as a negative control on the ELISA plates. Similarly, a sample of pooled sera from three 24-day-old birds that were vaccinated at day 1 and 14 post-hatching during a separate experiment was used as a positive control. Because of the high number of serum samples, multiple 96-well plates were used for the ELISA procedure. Cross-plate standardisation was performed by calculating the sample-to-positive ratio, using the following formula:

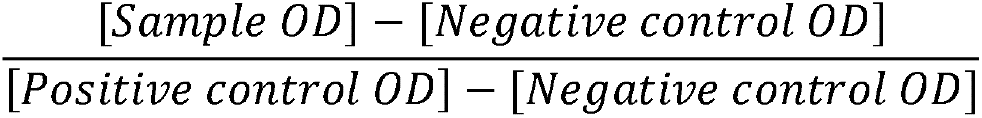

### 2.5. qRT-PCR

Total RNA from buccal and cloacal swab samples from H9N2 infected birds was extracted using the QIAamp Virus BioRobot MDx Kit (Qiagen) on a Biorobot Universal (Qiagen). Total RNA concentration was then measured using the NanoPhotometer NP80 (IMPLEN GmbH) to standardise across samples. A quantitative real-time PCR (qRT-PCR) using H9N2 matrix (M) gene primers was performed on samples using an M gene of known concentration (8.5 x 10^7^ copies/µl) as a standard, which was previously generated within the AIV group at the Pirbright Institute. For the qRT-PCR reactions, the Superscript III Platinum One-Step qRT-PCR Kit (Invitrogen) was used following the manufacturer’s instructions on a Quantstudio 5 Real-Time PCR System (Thermo Fisher Scientific). The results were then analysed using the Quantstudio 5 software (Thermo Fisher Scientific).

### 2.6. RNA extraction from tissues

RNA isolation from chicken tissue samples were carried as described in our previous work (18). 15 mg of tissue samples were transferred to tubes containing 600 µl of ice cold RLT lysis buffer (Qiagen) and 100 µl of 0.2 mm silica beads. Tissues were homogenised in a Mini-Beadbeater-24 (BioSpec) using 5 cycles of 1 minute each. Homogenate was then cooled on ice for 30 seconds, and RNA was extracted using the RNeasy Mini kit (Qiagen). On-column genomic DNA digestion was carried out using the RNase-Free DNase Set (Qiagen). Extracted RNA was eluted in 40 µl of nuclease-free water and RNA quality and integrity were assessed using an RNA ScreenTape (Agilent Technologies) and a 4200 TapeStation (Agilent Technologies), respectively. Samples were immediately stored at - 80°C.

### 2.7. cDNA generation and 5’RACE PCR

5’RACE-ready cDNA was generated using the SMARTer kit (Takara) according to the manufacturer’s instructions, and then 5’RACE PCR was carried out to amplify B cell receptor genes. 5’RACE PCR involved the use of 7-bp barcoded forward primers for immunoglobulin C_μ_ (NNNNNNNCACAGAACCAACGGGAAG), and immunoglobulin C_γ_ (NNNNNNNCGGAACAACAGGCGGATAG). The 5’RACE PCR was carried out in accordance with our previous protocol (18). This involved, universal SMARTer kit reverse primers that were specific to the common 5’ adapter added during 5’RACE cDNA synthesis. 5’RACE PCR reaction mixes contained: 5 µl of Phusion 5X Buffer (New England Biolabs), 0.5 µl of 10 mM dNTP, 0.5 µl of 10 µM UPA-short primer, 0.5 µl of 2 µM UPA-long and 0.25 µl Phusion Hot Start Flex DNA Polymerase (New England Biolabs) were added to 15.25 µl nuclease-free water, for a total of 22 µl volume. To this, 0.5 µl of the 10 µM gene-specific 7bp-barcoded primer and 2.5 µl of cDNA were added. The individual 25 µl volume 5’RACE PCR reactions were then carried out in 96-well plates using the thermocycler program recommended by the 5’RACE kit (Takara), with 35 cycles of gene-specific amplification with an annealing temperature of 60°C. Barcoded PCR products were pooled and subjected to electrophoresis on a 1.4% agarose in 45 mM Tris-Borate/ 1mM EDTA (TBE) buffer gel containing 1:10,000 SYBR green (Sigma-Aldrich) at 120 V for 35 minutes. The bands of the expected lengths were gel extracted and purified using the QIAquick Gel Extraction Kit (Qiagen).

### 2.8. DNA library preparation and sequencing

Pooled barcoded PCR products were used to generate DNA libraries using the NEBNext Ultra II DNA Library Prep Kit for Illumina (New England Biolabs). A NEBNext Library Quant Kit for Illumina (New England Biolabs) and a D1000 DNA tape (Agilent Technologies) for the 4200 TapeStation (Agilent Technologies) were used to assess the quantity and quality of the DNA libraries. Library preparation and sequencing was carried out using an Illumina MiSeq platform by the Bioinformatics, Sequencing & Proteomics group at the Pirbright Institute.

### 2.9. Repertoire sequence data processing and analysis

An in-house python package (available on GitHub: https://github.com/sgp79/reptools) was used to process the raw sequencing data using the same workflow established previously (18,19). Briefly, the software identifies the V and J gene IDs to the sequences by BLAST (20). The algorithm then extracts the CDR3 sequences after a Smith-Waterman alignment (21), allowing for higher precision at the junctions (22). The output was then analysed using R (23).

### 2.10. Repertoire diversity

Patterns of clonal expansion with respect to tissue type and immunisation regime were investigated by analysing the diversity present within samples, as established previously (18). Briefly, the effective number of species, here equating the “effective number of clones” (D) can be calculated using the weighted abundance of unique reads in each sample (24,25). Diversity was measured using the following definitions: clonal richness (D0 – the total number of clones found regardless of abundance), the exponential of the Shannon entropy (D1 – defined here as “typical clones”), and the inverse Simpson concentration (D2 – defined here as “dominant clones”). Diversity measures were computed, thus incorporating the effect of clonal expansions at different levels. This was achieved using the iNEXT package (26) in R, and interpolation or extrapolation was carried out to standardise between samples at a value of 1,000 total reads.

### 2.11. Public and private clonal compartments

Public and private clonal compartments were analysed using previously established methods (18). Briefly, CDR3 nucleotide sequences were divided into private or public compartments depending on whether they were shared between individuals. The percent contribution to the private or public clonal compartment was estimated as a function of immunisation regime, tissue type, and clonal compartment (private or public). At first, all clones which were found in more than one individual were regarded as public, irrespective of the number of individuals which shared them. Subsequently, a new model was constructed where public clones were divided into multiple categories based on their presence across birds: rare publics (2 or more, up to 50% of birds), common publics (more than 50%, up to 90% of birds), and ubiquitous clones (more than 90% of birds).

## 3. Results

### 3.1. Confirmation of vaccination and/or infection status of birds

AIV reactive IgM and IgY antibody titres were measured by ELISA and HI assays prior to and during the study. This confirmed the vaccinated or infected status of birds as planned (Supplementary Figure 2). Buccal and cloacal swabs were collected from all bird groups daily until 8 days post infection (dpi). The buccal swabs confirmed the infection status (Supplementary Figure 3) via qRT-PCR for the M gene of the virus, while the cloacal swab samples did not exhibit any detectable levels of viral shedding (data not shown).

### 3.2. Productive rearrangement of IgM & IgY sequences

Sequencing of 5’RACE PCR products yielded a total of 2,295,729 IgM reads, and 2,165,701 IgY reads. Of these, ∼97.3% and ∼96.1% were productively rearranged (i.e. could yield a protein product free from premature stop codons or frameshifts), respectively. There was substantial heterogeneity in total reads obtained across individual samples (Supplementary Figures 4 & 5), with no consistent patterns being observed between immunisation regimes or tissue types. Almost all IgM samples had more than 1000 productive reads, except for the trachea of one bird from which 5 sequences were recovered, of which 3 were productively rearranged. This tissue sample was excluded from the analysis, due to the insufficient number of reads. Four IgY samples were also excluded from the proportion-based analyses as they had ≤500 productively rearranged reads. This was done to minimise the influence of potentially dominant clones on the repertoire composition of the samples in question. The immunisation regime groups affected by the removal of these samples had at least n=4 samples per tissue, which is comparable to the samples of the groups with the lowest number of birds (i.e. the single vaccination and single vaccination and infection).

### 3.3. IgM clonal homeostasis

To determine the effect of immunisation regime on the proportion of IgM clonotypes, we generated IgM clonal homeostasis plots for the spleen, bursa, and trachea for each immunisation regime (Figure 1). Although there was no apparent regime-specific effect, apart from relatively reduced clonal expansion in the spleen of double vaccinated birds, there were more clear differences between the tissues, whereby the trachea displayed the highest proportion of expanded clones, and the bursa the lowest.

**Figure 1:**
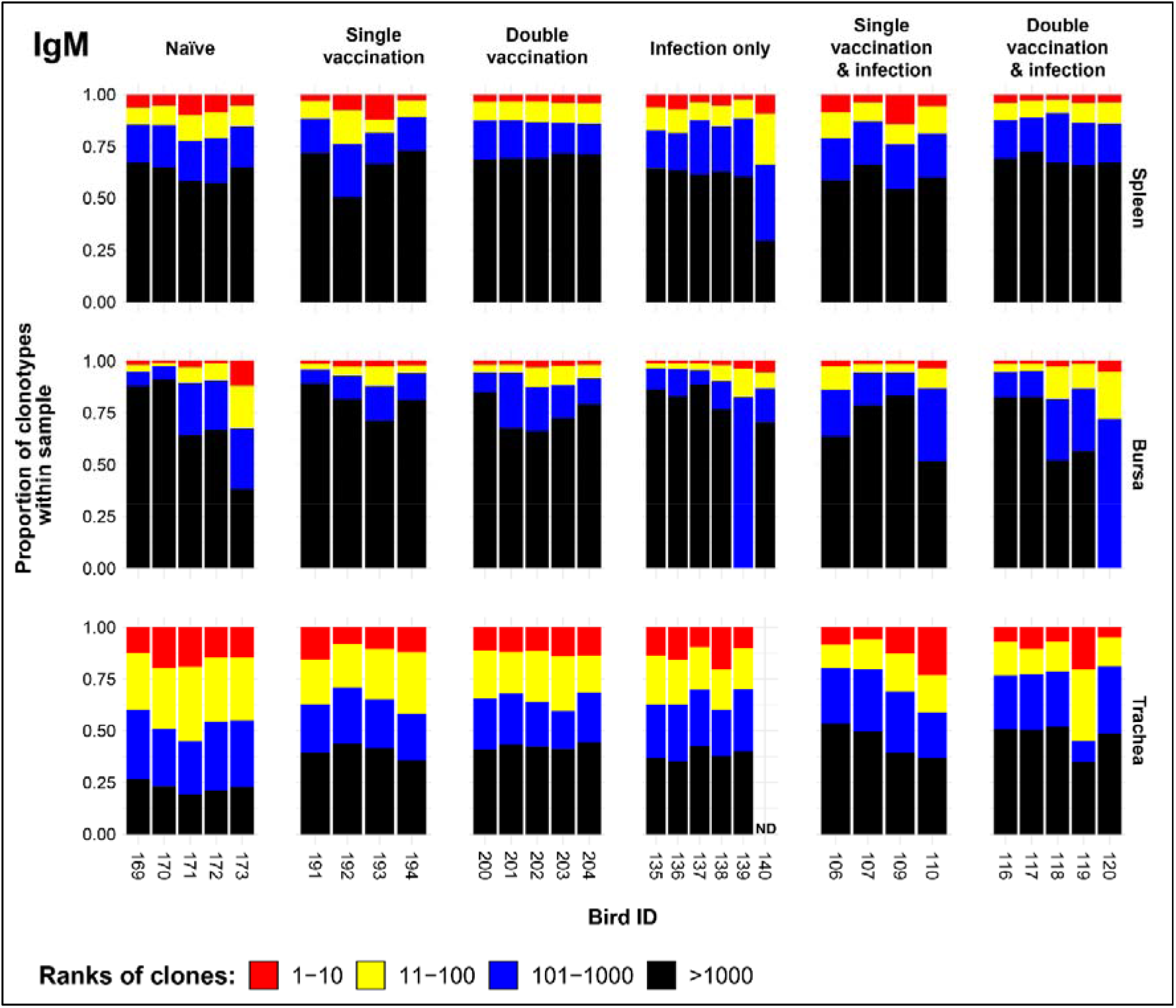
IgM clonal homeostasis plots of individual tissue samples. Bird numbers are displayed on the x axis and individuals are grouped based on immunisation status which is illustrated above each panel. Clones were ranked based on their abundance into four categories: first 10 most abundant (red), from 11-100 (yellow), 101-1000 (blue), and above 1000 (black) in terms of total abundance within each sample. The proportions of clonotypes are displayed on the y axis.

### 3.4. IgM repertoire diversity

We next investigated the IgM clonal diversity across tissues and immunisation regimes (Figure 2). The bursa showed no differences in clonal diversity. The spleen showed differences between naïve birds and the double vaccinated, and the double vaccinated and infected dominant clones (D2) only, which both exhibited significantly higher levels of diversity. In contrast, the trachea showed significantly higher levels of diversity in all groups in both clonal richness (D0) and typical clones (D1), when compared to the naïve group. However, only the double vaccinated and infected group showed significantly higher diversity to the naïve birds when looking at the dominant clones (D2).

**Figure 2:**
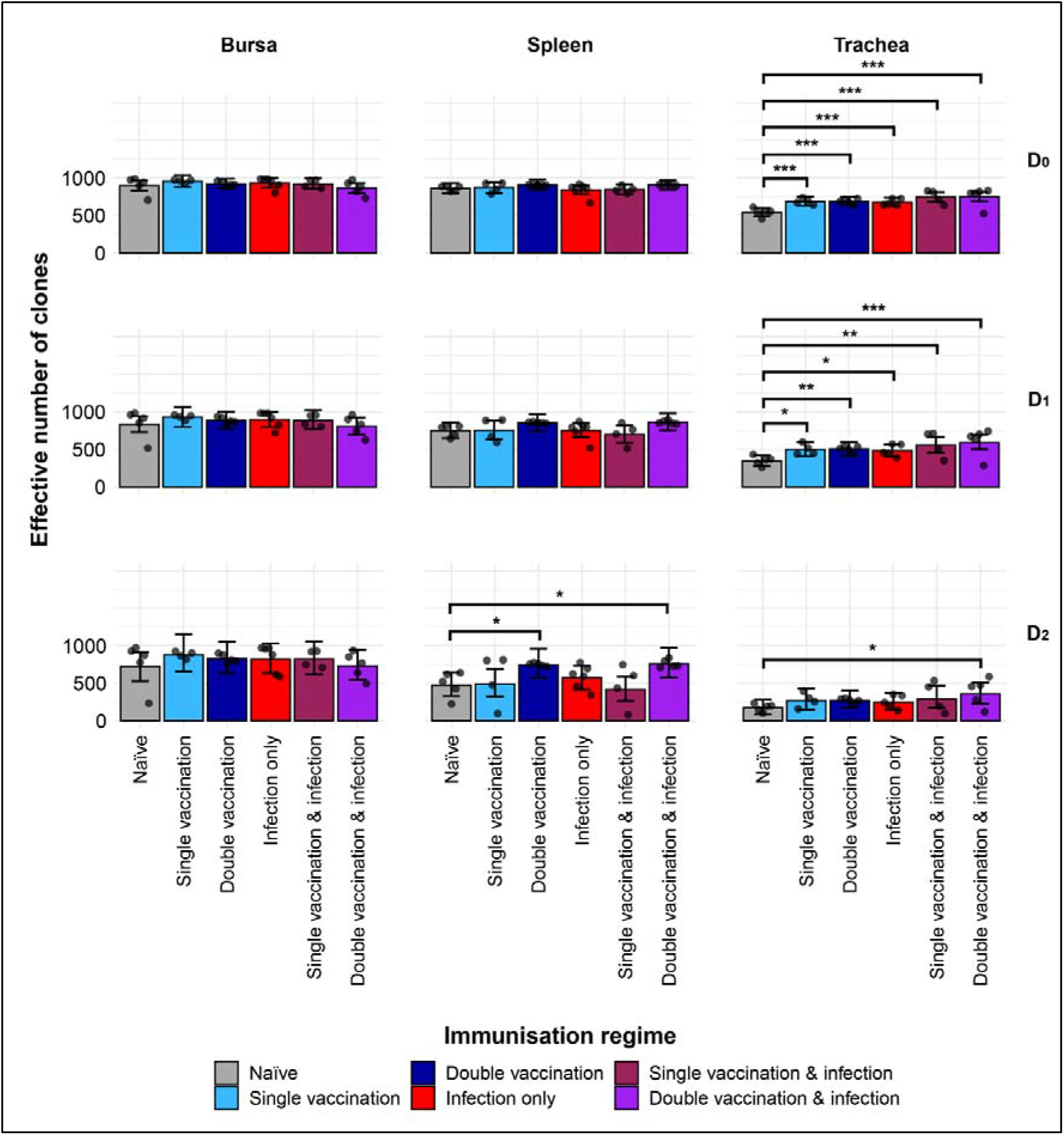
IgM clonal diversity within samples. Different rows show the Hill numbers corresponding to clonal richness (D0), the “typical” clones (D1) and the “dominant” clones (D2) in a theoretical sample of 1000 sequences. Immunisation regimes are colour coded and displayed on the x axes. Dots represent individual bird observations of the effective number of species calculated in each tissue for the corresponding H values. Error bars show the 95% bootstrap confidence intervals for the point estimates generated from 1000 simulations of the model. Statistically significant differences between the model estimates are depicted above the plots based on their corresponding p-values: * = p < 0.05; ** = p < 0.01; *** = p < 0.001.

### 3.5. Public and private IgM clonal compartments

The partitioning of IgM sequences into private and public clones (at the nucleotide level) reveals effects of both tissue type and immunisation regime on the composition of the repertoires (Figure 3). The bursal tissue samples of all groups exhibited significantly more private than public clones, with the notable exception of the double vaccination and infection group, which had comparable proportions of public and private clones. By contrast, the tracheal samples of groups exhibited significantly higher proportions of public clones than private, with the former reaching more than 75% of the total clones in some individuals. In the spleen, the infection only group had significantly more private clones than public. At the same time, the single vaccination and infection group exhibited more public than private clones in the spleen. The splenic samples from other groups had comparable levels of private and public clones.

**Figure 3:**
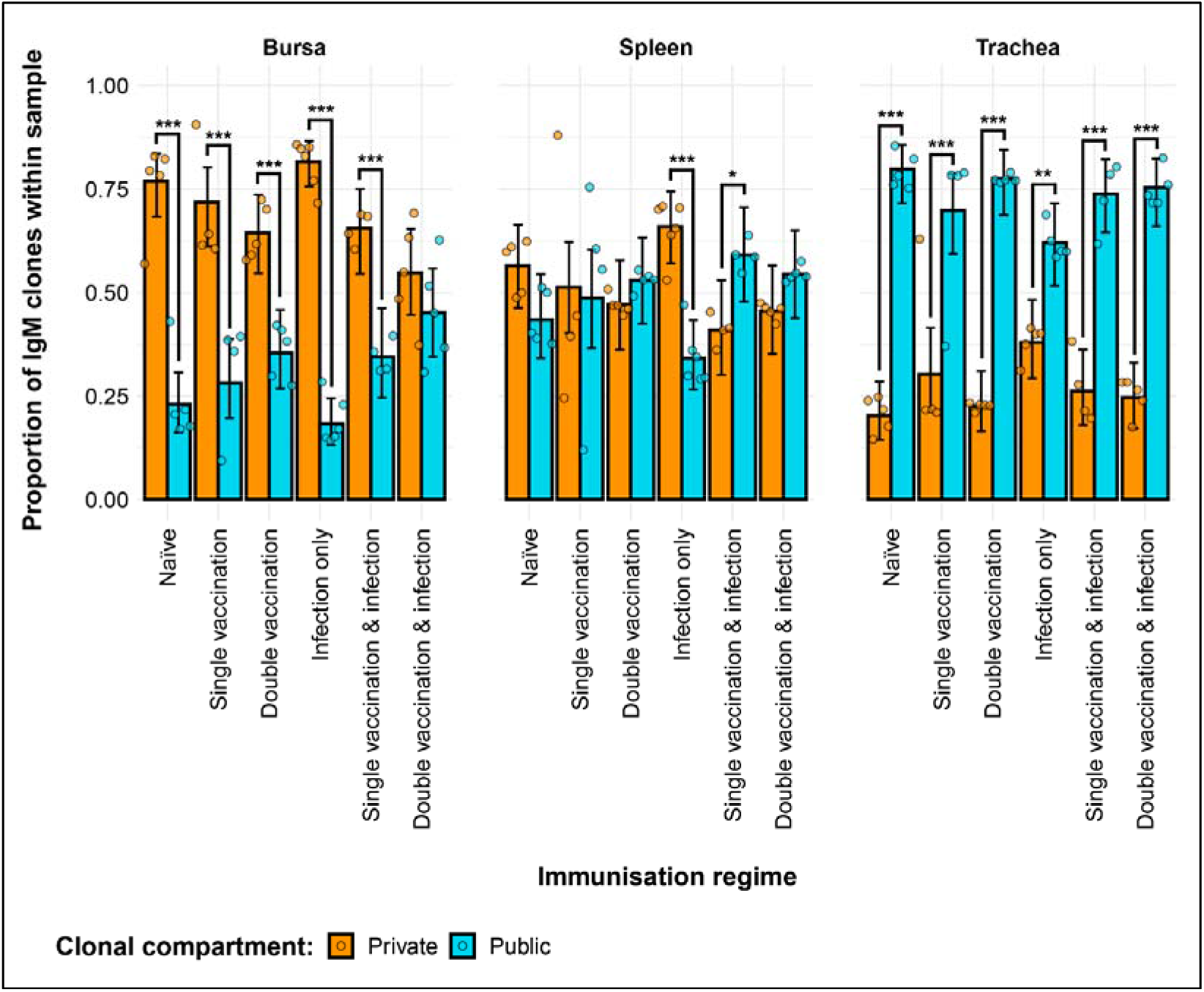
Differences between the IgM public and private compartments under different H9N2 immunisation regimes based on clone CDR3 nucleotide structure. Private (individual-restricted) clones are shown in orange. Public clones (shared between more than two individuals) and are shown in light blue. Dots represent individual bird observations of public and private clonal compartments. Error bars represent 95% bootstrap confidence intervals for the point estimates generated from 1000 simulations of the model. Statistically significant differences between the model estimates are depicted above the plots based on their corresponding p-values: * = p < 0.05; ** = p < 0.01; *** = p < 0.001.

Differences between the groups were also evident regarding the contribution of private and public clones to the repertoire (Figure 4). In the bursa, the double vaccination regimes exhibited lower proportions of private clones and higher proportions of public clones. In the trachea, the only immunisation regime that exhibited a difference to the naïve birds was the infection only group, showing significantly higher proportions of private clones and lower proportions of public clones.

**Figure 4:**
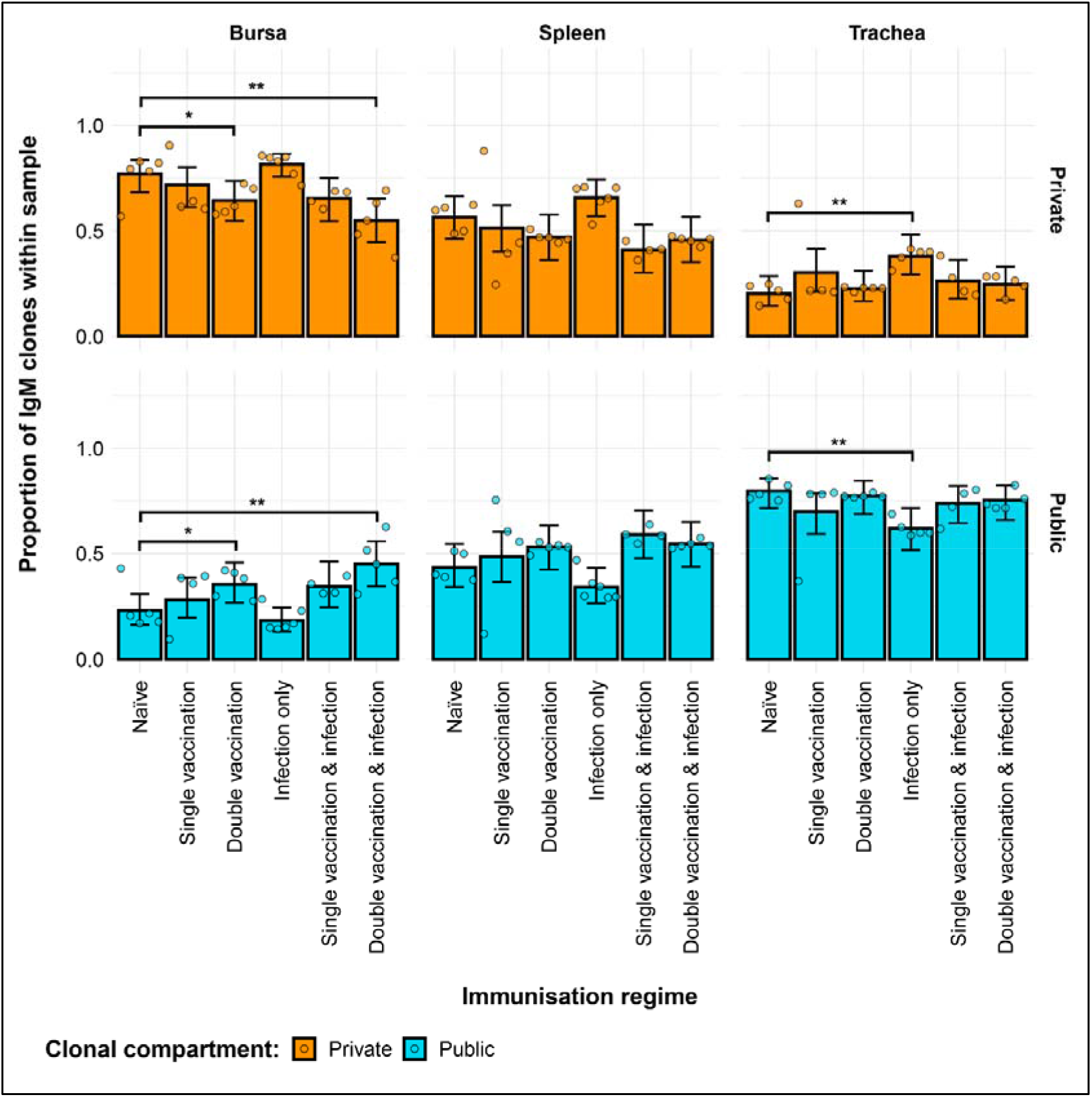
Differences within the IgM public and private compartments under different H9N2 immunisation regimes based on clone CDR3 nucleotide structure. Private (individual-restricted) clones are shown in orange. Public clones (shared between more than two individuals) and are shown in light blue. Dots represent individual bird observations of public and private clonal compartments. Error bars represent 95% bootstrap confidence intervals for the point estimates generated from 1000 simulations of the model. Statistically significant differences between the model estimates are depicted above the plots based on their corresponding p-values: = p < 0.05; ** = p < 0.01; *** = p < 0.001.

The partitioning of public IgM clones based on their degree of clonal sharing reveals additional effects between the samples (Figure 5). Notably, the most substantial component of the total public repertoire is comprised of rare public clones. As such, the differences to the naïve birds regarding the proportion of total public clones were also displayed by the rare public compartment of the immunisation regimes across tissues. When looking at the private clones, the differences to the naïve group in the trachea and bursa were also consistent with the ones revealed by the model without the partitioning of the public compartment. However, in the spleen, the single vaccination and infection group now exhibited significantly lower proportions of public clones to the naïve.

**Figure 5:**
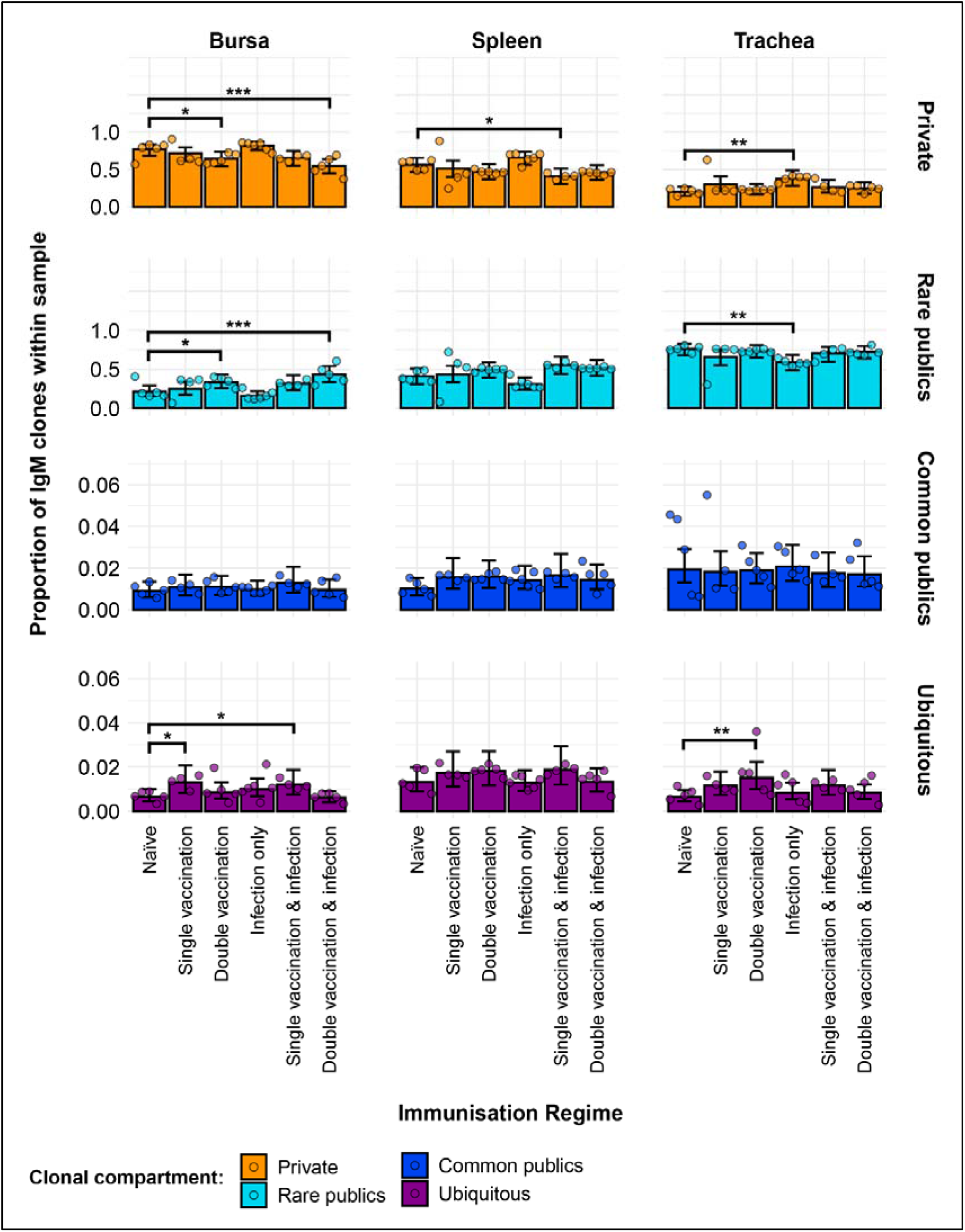
Model estimates of IgM clone CDR3 nucleotide private and public compartments based on different levels of clonal sharing. Private (individual- restricted) clones are shown in orange. Rare publics (shared between 2 or more than 2 individuals up to 50%) and are shown in light blue. Common publics (shared between more than 50% and up to 90% birds) are shown in dark blue. Ubiquitous publics (found in 90% or more of the birds which were incorporated in the analysis) are shown in purple. Dots represent individual bird observations of private and distinct public clonal compartments. Error bars represent 95% bootstrap confidence intervals for the point estimates generated from 1000 simulations of the model. Statistically significant differences between the model estimates are depicted above the plots based on their corresponding p-values: * = p < 0.05; ** = p < 0.01; *** = p < 0.001.

Regarding the more frequently shared IgM clones, both the common public and the ubiquitous clonal compartments were at relatively low levels, each generally amounting to less than 5% of the repertoire of individual birds. There were no significant differences between the IgM repertoires of the immunisation regimes regarding the common clones, but some were evident in the ubiquitous clonal compartments. In the bursa, the single vaccination and the single vaccination and infection groups showed significantly higher levels than the naïve birds. In the trachea, the double vaccinated group had significantly higher proportions of ubiquitous clones than the naïve chickens.

### 3.6. IgM public repertoires restricted to immunisation regimes

Only 3 IgM clones which were present and expanded across multiple individuals were found restricted to the infected birds, and unexpanded or absent in the uninfected birds (Figure 6). Of these, one clone (CDR3: CAKSTAGTCWYDDAGSID) was present only in birds from the vaccination and infection treatments. This clone was expanded in the spleen and tracheas of two single vaccinated and infected birds. Additionally, one of the single vaccination and infection birds also showed an expansion of this clone in the bursa, whereas the clone was not detected in the other bird. This clone was also identified in the bursa of a bird from the double vaccinated and infected group, but it is not expanded.

**Figure 6:**
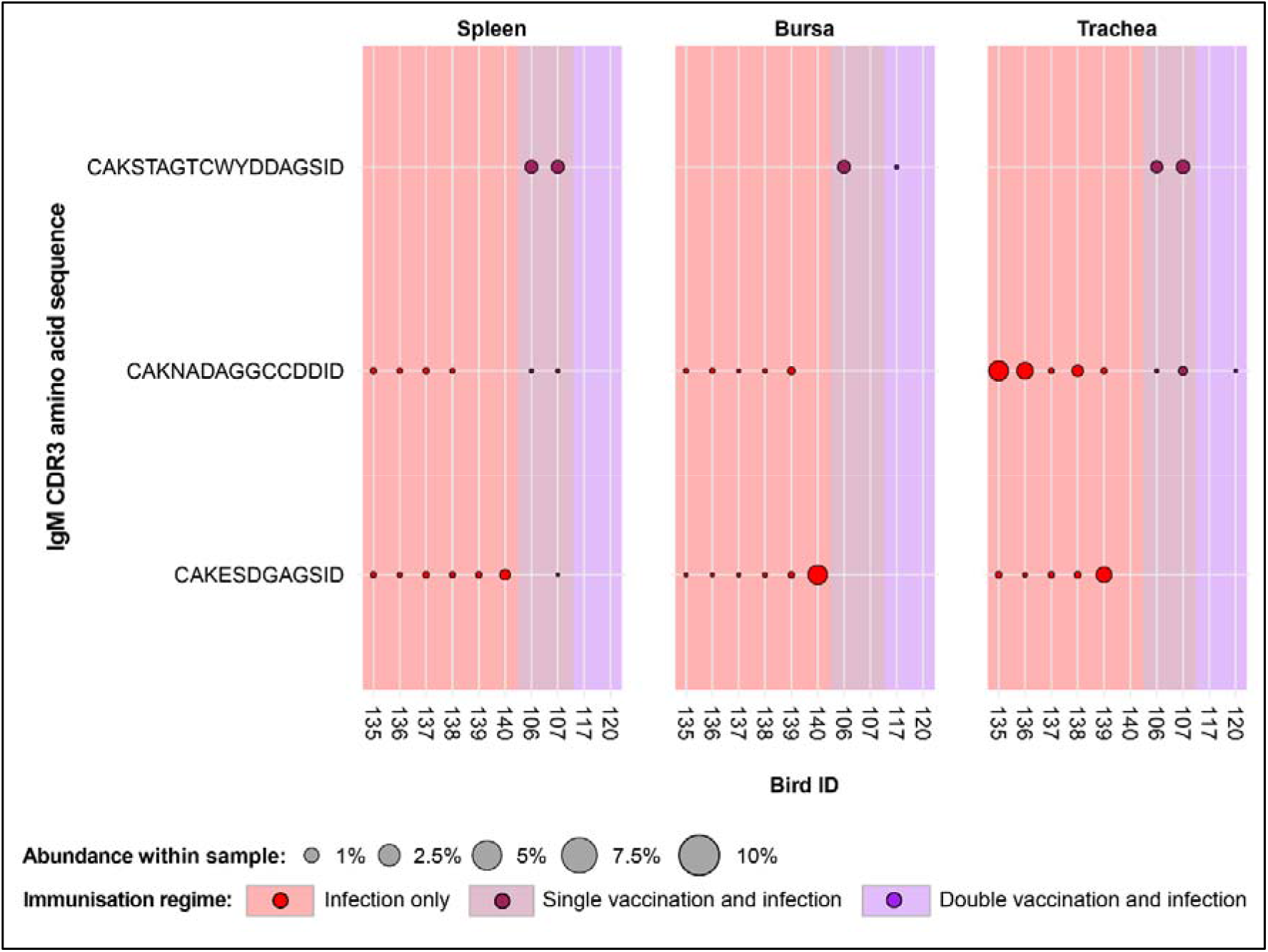
IgM Clonal expansions in the restricted repertoire of infected birds. Circles indicate the presence of clones with a specific CDR3 amino acid sequence and are proportional to the abundance within each bird’s tissue clonal compartment. The plot shows only the clones which are expanded in the infected treatment groups (at or above 0.5% and at less than 0.5% in any of the uninfected). Background and circle colours indicate the immunisation regime identity: red, infection only; dark red, single vaccination and infection; purple, double vaccination and infection.

The remaining two public IgM clones restricted to the infected groups were identified in both infection-only birds and single vaccinated and infected individuals, although expansions were present only in infection-only birds. One clone (CDR3: CAKESDGAGSID) was present in almost all the analysed tissues of the individuals belonging to the infection-only group, and in one splenic sample of a single vaccinated bird. The expansions of this IgM clone were identified in the trachea of one bird, and bursa of another. Interestingly, the clone was not identified in the trachea of the bird that exhibited an expansion in the bursa. The other clone (CDR3: CAKNADAGGCCDDID) was found in both the infection only and single vaccination and infections groups but only showed expansions in 3 tracheal samples of the infection-only birds.

Expansion patterns were found when observing IgM clones in the uninfected treatment groups (Figure 7a). Expansions were only observed in the vaccinated groups and not in the naïve birds. Of these, three clones were found expanded in the single vaccination, and one clone in the double vaccination treatment. In the former group, all 3 clones are present in all analysed tissues of 3 birds, although their patterns of expansion differ. One clone (CDR3: CAKGSGCCGSRGRTAGTID) was found at comparable proportions (∼1%) in the tissues of the 3 birds. Another clone (CDR3: CAKSSYECAYDCWGYAGSID) also displayed expansions across all tissues of the three single vaccinated chickens, particularly in the spleen of one bird, where it reached close to 10% of the repertoire. The remaining one (CDR3: CAKSYGGNWGGFIEDID) was only expanded in the spleen and trachea of one single vaccinated bird, but still present in the bursa and in all the tissues of the remaining two birds. The clone that was expanded in the double vaccinated birds (CDR3: CAKESSSAVSID) was present in the five individuals of the group in all tissues, except for one bursal sample. The identified expansions were in the tracheas of two birds, reaching ∼0.5% and ∼2% of the repertoire, respectively. When considering the public IgM expansions restricted to the vaccination groups (including under infection settings), only one clone (CDR3: CAKESSYADSID) is expanded in both infected and uninfected vaccinated groups (Figure 7b). In the double vaccinated treatment group, this IgM clone was expanded in all splenic and tracheal samples. Furthermore, this clone was present in all bursal tissues of the double vaccinated birds, but only expanded in one. This clone was also identified in the bursa and trachea of one single vaccinated bird, although it was unexpanded. However, in the double vaccination and infection group, the clone is present at unexpanded levels in the spleen, and in 3 bursal samples. In the trachea, all birds of the double vaccinated and infected possess this clone, with one expansion being present.

**Figure 7:**
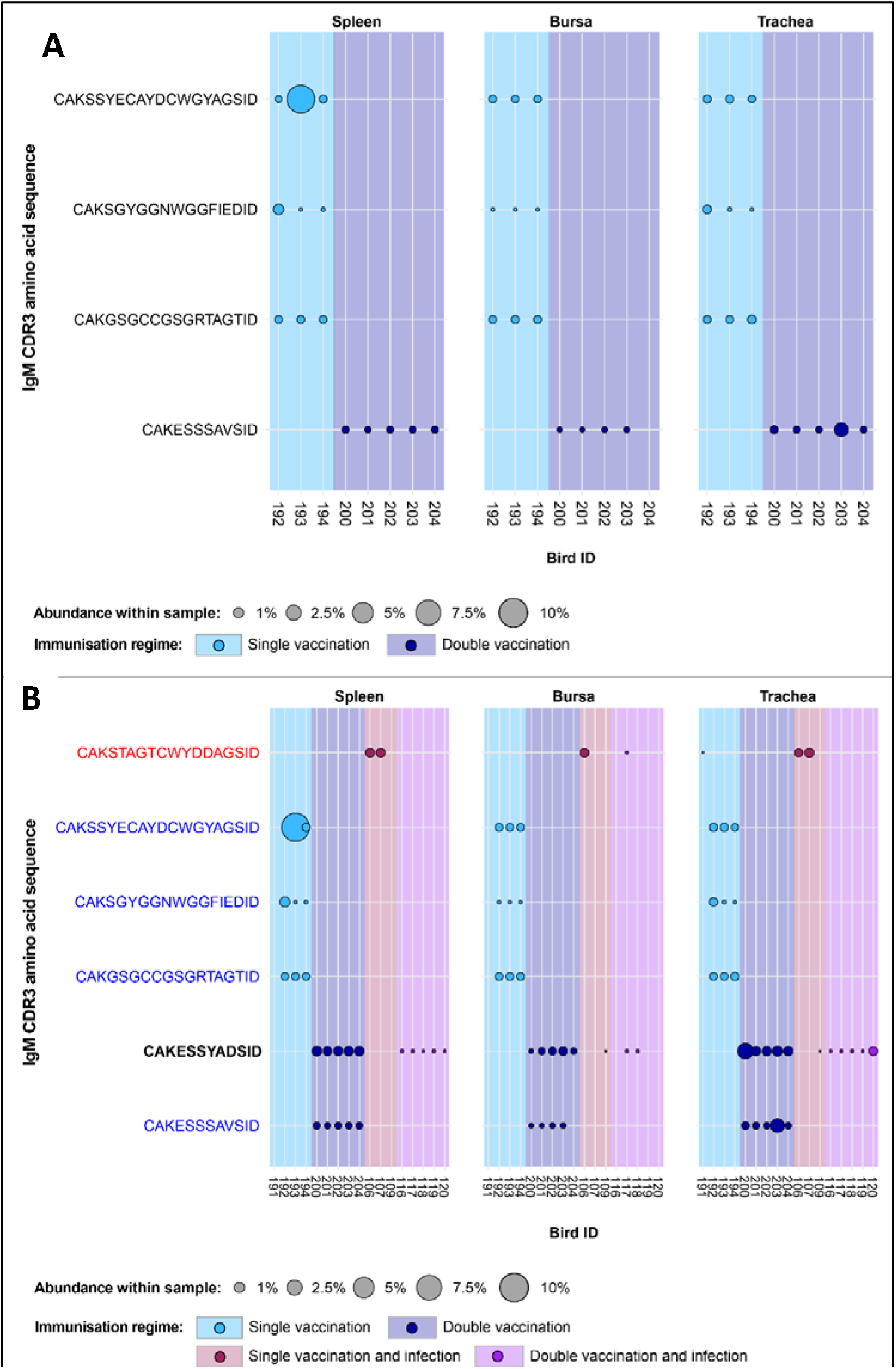
IgM Clonal expansions in the restricted repertoire of uninfected birds and of vaccinated birds. Circles indicate the presence of clones with a specific CDR3 amino acid sequence and are proportional to the abundance within each bird’s tissue clonal compartment. (**A**) The plot shows only the clones which are expanded in the uninfected treatment groups (at or above 0.5% and at less than 0.5% in any of the infected). (**B**) The plot shows only the clones which are expanded in the vaccinated treatment groups (at or above 0.5% and at less than 0.5% in any of the naïve or infection only groups). CDR3 amino acid sequences which coincide with clones restricted to infected and uninfected birds are shown in red and blue, respectively. Background and circle colours indicate the immunisation regime identity: light blue - single vaccination; dark blue - double vaccination, dark red - single vaccination and infection; purple - double vaccination and infection.

### 3.7. IgY clonal homeostasis

The patterns of clonal homeostasis illustrated the influence of tissue type and treatment group on the IgY repertoire (Figure 8). The tracheal samples generally exhibit the highest levels of expansion followed by the spleen and then the bursa, irrespective of the immunisation regime. Heterogeneity between samples is pronounced, indicating that the influence of the individual is important. In the bursa and spleen, the clonal homeostasis patterns between the groups are generally comparable to one another. The trachea exhibits the highest levels of IgY clonal expansions, most notably in the uninfected groups.

**Figure 8:**
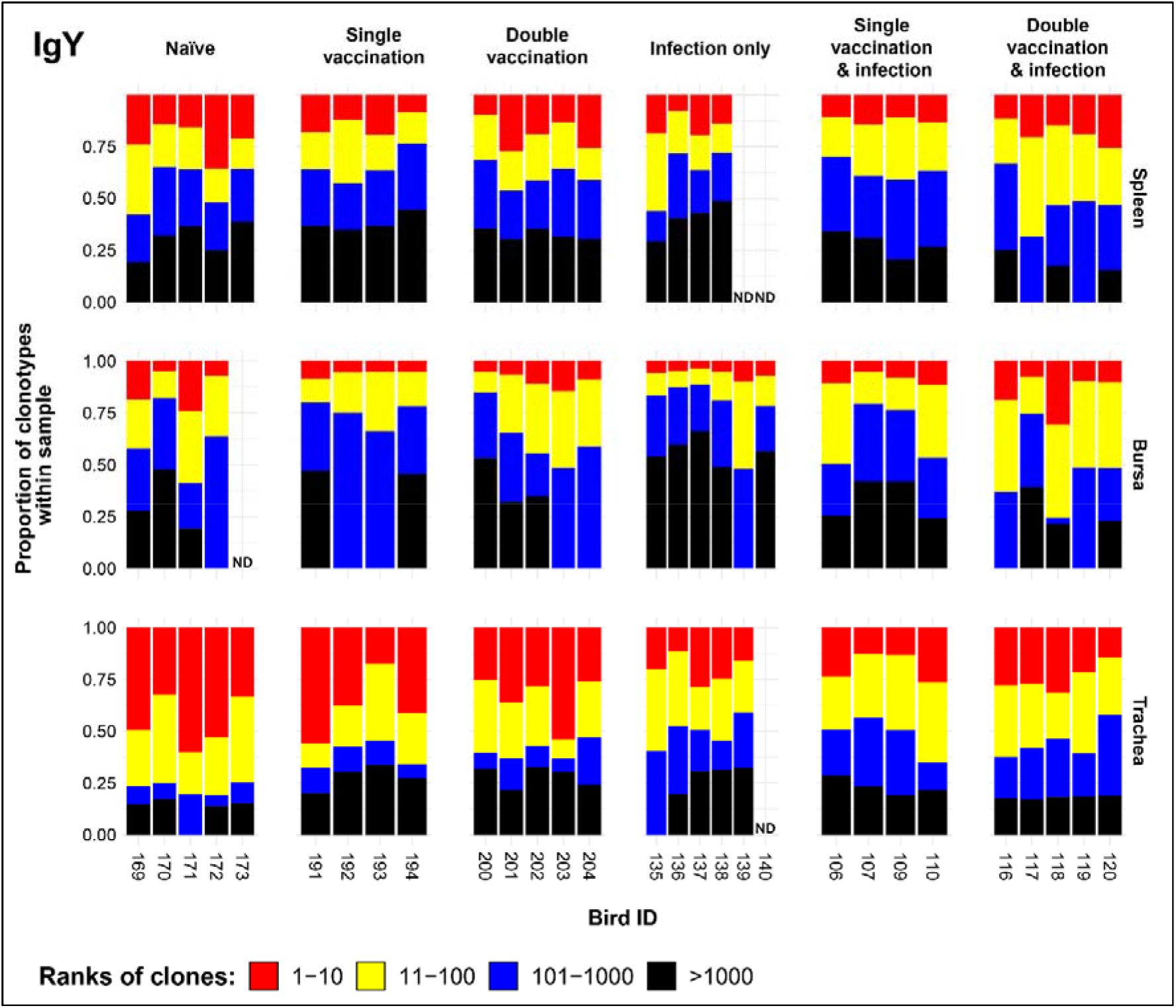
IgY clonal homeostasis plots of individual tissue samples. Bird numbers displayed on the x axis and individuals are grouped based on the corresponding immunisation status which is illustrated above each panel. Clones were ranked based on their abundance into four categories: first 10 most abundant (red), from 11-100 (yellow), 101-1000 (blue), and above 1000 (black) in terms of total abundance within each sample. The proportions of clonotypes are displayed on the y axis. Samples which were removed due to a low read number are displayed as having no data (ND) at the corresponding locations.

### 3.8. IgY repertoire diversity

The IgY diversity patterns provided further information on the IgY clonal landscapes within the analysed tissues (Figure 9). In the bursa, the groups that received one immunisation, through either vaccination or infection are significantly more diverse in terms of clonal richness (D0) than the naïve treatment. These differences are maintained when considering the typical clones (D1) or the dominant clones (D2). The diversity of the splenic samples for the immunisation regimes and the naïve birds showed no significant difference in either clonal richness or typical clones. However, when looking at the dominant clones, the single vaccination and infection group has significantly more diverse repertoires than the naïve group. The single vaccination treatment also seems to be more diverse than the naive, although this difference was not statistically significant. In the trachea, all immunised groups are statistically more diverse in terms of both IgY clonal richness and typical IgY clones than the naïve group. When considering the dominant clones, however, the infection only and the single vaccination and infection groups have significantly higher levels of diversity than the naïve birds in the trachea.

**Figure 9:**
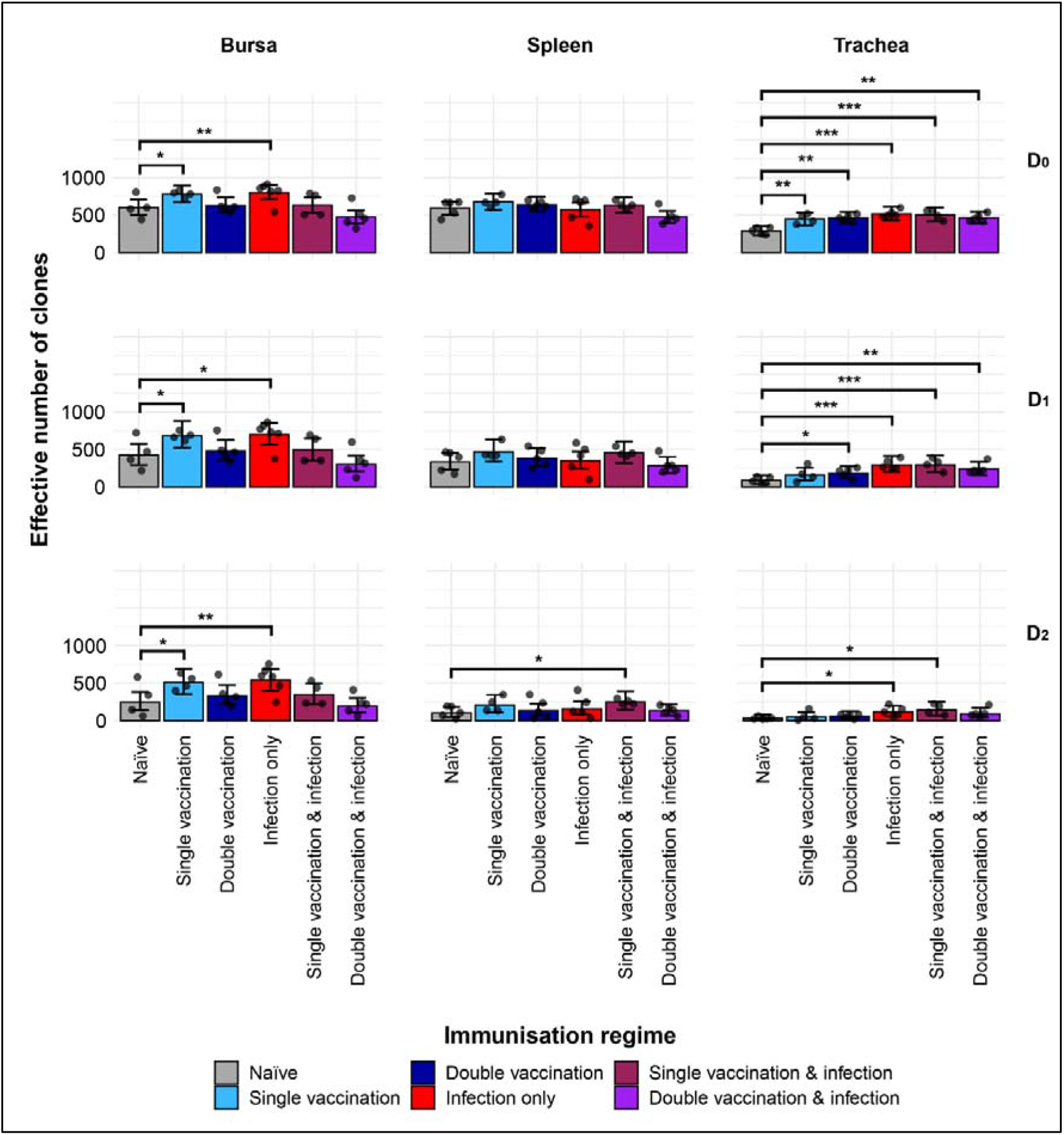
IgY clonal diversity within samples. Different rows show the Hill numbers corresponding to clonal richness (D0), the “typical” clones (D1) and the “dominant” clones (D2) in a theoretical sample of 1000 sequences. Immunisation regimes are colour coded and displayed on the x axes. Dots represent individual bird observations of the effective number of species calculated in each tissue for the corresponding H values. Error bars show the 95% bootstrap confidence intervals for the point estimates generated from 1000 simulations of the model. Statistically significant differences between the model estimates are depicted above the plots based on their corresponding p-values: * = p < 0.05; ** = p < 0.001. *** = p < 0.01;

### 3.9. Public and private IgY clonal compartments

The IgY repertoires of the H9N2 immunisation regimes exhibited significant differences in terms of their private and public clones (Figure 10). In the bursa, the naïve, single vaccination, and infection only groups showed significantly higher contributions of private clones than public clones to the repertoire. The opposite pattern was observed for the double vaccination treatment which exhibited significantly higher contributions of public clones as opposed to private clones. The vaccinated and infected groups showed similar contributions in terms of public and private clones to the repertoire. In the spleen, the double vaccinated group and the single vaccination and infected group exhibited significantly higher proportions of public rather than private clones. Lastly, all tracheal samples of the immunisation regimes had significantly higher proportions of public clones as opposed to private, which reached more than 50% of the repertoire on most occasions, with public clones exceeding 80% of the total clones.

**Figure 10:**
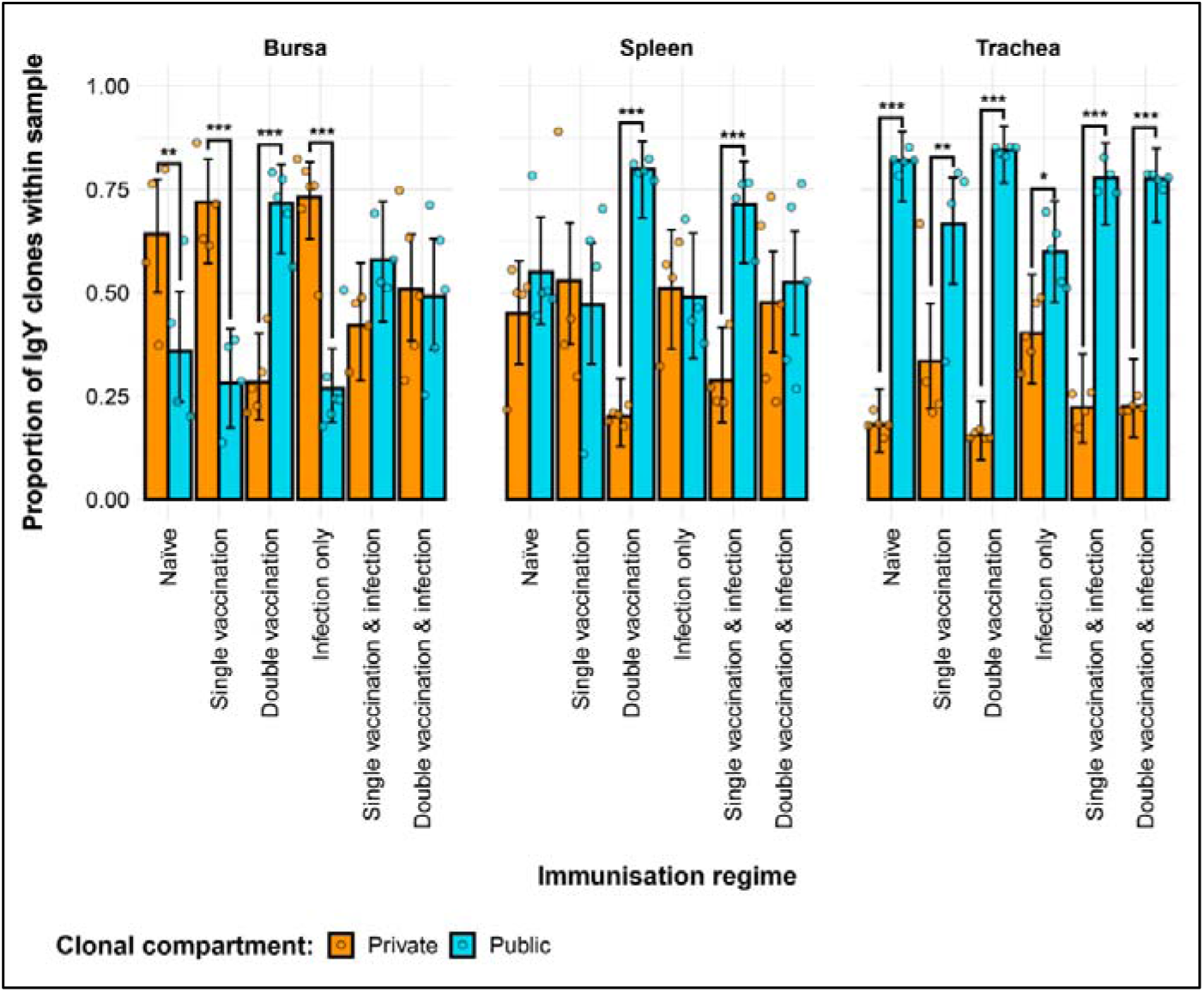
Differences between the IgY public and private compartments under different H9N2 immunisation regimes based on clone CDR3 nucleotide structure. Private (individual-restricted) clones are shown in orange. Public clones (shared between more than two individuals) and are shown in light blue. Dots represent individual bird observations of public and private clonal compartments. Error bars represent 95% bootstrap confidence intervals for the point estimates generated from 1000 simulations of the model. Statistically significant differences between the model estimates are depicted above the plots based on their corresponding p-values: * = p < 0.05; ** = p < 0.01; *** = p < 0.001.

The private and public clonal compartments of the immunised groups also differed in terms of their relative sizes in the repertoire when compared to the naïve birds (Figure 11). In the bursa, the double vaccination regime and the single vaccination and infection birds had significantly higher proportions of public clones and lower levels of private clones than the naïve group. In the spleen, only the double vaccinated birds exhibited a significant difference to the naïve, having higher levels of public clones and lower levels of private clones. Lastly, the tracheal samples of the singly immunised birds, either through vaccination or infection, exhibited higher proportions of private clones and lower proportions of public clones than the naïve birds.

**Figure 11:**
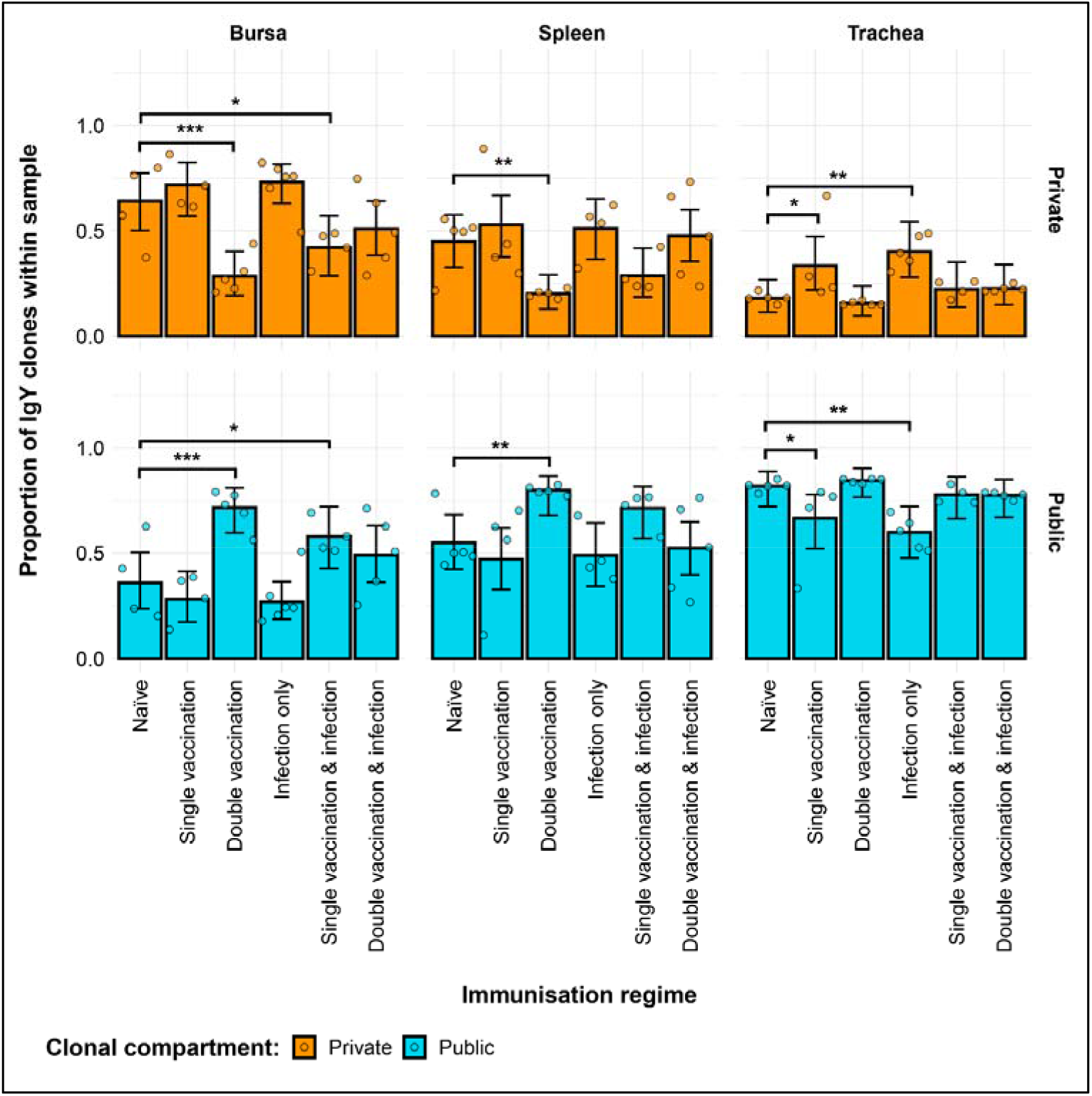
Differences within the IgY public and private compartments under different H9N2 immunisation regimes based on clone CDR3 nucleotide structure. Private (individual-restricted) clones are shown in orange. Public clones (shared between more than two individuals) and are shown in light blue. Dots represent individual bird observations of public and private clonal compartments. Error bars represent 95% bootstrap confidence intervals for the point estimates generated from 1000 simulations of the model. Statistically significant differences between the model estimates are depicted above the plots based on their corresponding p-values: * = p < 0.05; ** = p < 0.01; *** = p < 0.001.

Further partitioning of the public clones into categories based on the degree of clonal sharing between individuals revealed several patterns in terms of tissue and group-specific differences (Figure 12). Most public clones are rare publics (shared between 2 and up to 50% of all birds), irrespective of tissue type or immunisation regime. However, some of the previously observed differences between the groups were masked when partitioning the public clones through this model. As such, in the bursa and spleen, the single vaccinated treatments were not deemed to be significantly different anymore in terms of private clones to the naïve. The other previously described differences in terms of private clones remained significant, and were also mirrored in the rare public compartment, with the exception of the aforementioned samples of the single vaccination groups. Other significant differences between the immunised groups and the naïve birds were revealed when considering the common public clones (shared by ≥50% and <90% of birds) and the ubiquitous clones (found in ≥90% of individuals). In the bursa, the double vaccination and infection birds had significantly lower common public clones than the naïve, whilst the other groups do not exhibit any differences. In the trachea, all three infected groups, irrespective of vaccination status showed significantly higher proportions of common publics than the naïve group. No differences between the groups were observed in the spleen. By contrast, when considering the ubiquitous clones, there were no differences between the groups in either the trachea or the bursa, but one difference was apparent in the spleen. There, the ubiquitous clones of the double vaccination treatment were at statistically higher levels than the naïve group. Taken together, these results reveal interesting effects of both tissue type and H9N2 immunisation regime on the private and public clonal compartments of the birds.

**Figure 12:**
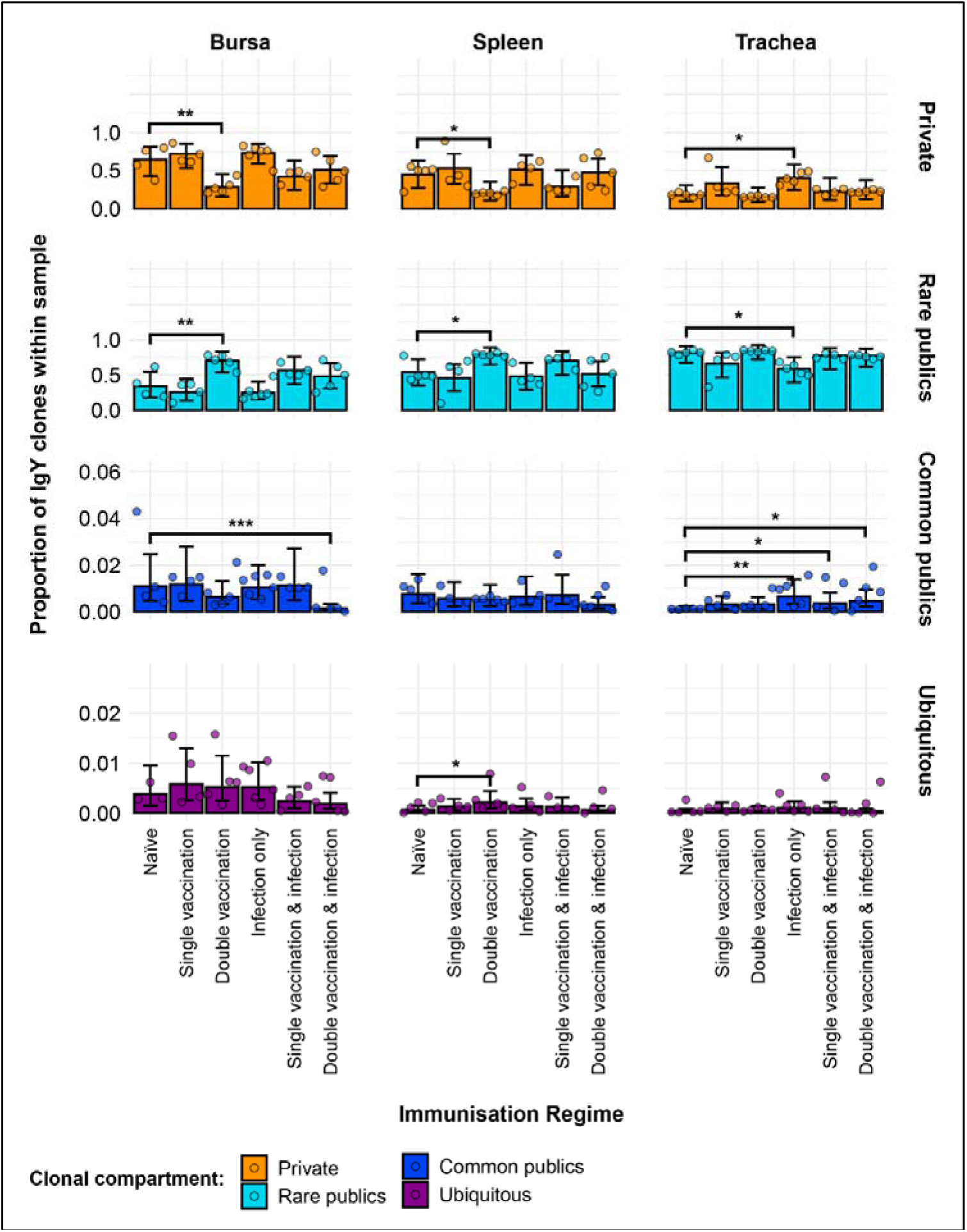
Model estimates of IgY clone CDR3 nucleotide private and public compartments based on different levels of clonal sharing. Private (individual- restricted) clones are shown in orange. Rare publics (shared between 2 or more than 2 individuals up to 50%) and are shown in light blue. Common publics (shared between more than 50% and up to 90% birds) are shown in dark blue. Ubiquitous publics (found in 90% or more of the birds which were incorporated in the analysis) are shown in purple. Dots represent individual bird observations of private and distinct public clonal compartments. Error bars represent 95% bootstrap confidence intervals for the point estimates generated from 1000 simulations of the model. Statistically significant differences between the model estimates are depicted above the plots based on their corresponding p-values: * = p < 0.05; ** = p < 0.01; *** = p < 0.001.

### 3.10. IgY public repertoires restricted to immunisation regimes

A total of 28 IgY public CDR3 amino acid clones were expanded in birds belonging to the infected groups and found absent or unexpanded in the uninfected groups (Figure 13). Of these, one CDR3 amino acid clone (CTKCAYSWCAAGSID) was comprised of two unique clonal lineages with different CDR3 nucleotide sequences. Although only showing expansions in the trachea and bursa of one double vaccinated and infected bird, it was found present at unexpanded levels in all the individuals of the group across multiple tissues and in the bursa of a single vaccination and infection bird. The majority of the other infection-restricted public clones are also only expanded in one individual and generally only in one tissue, in spite of them being shared across multiple birds. Only one clone (CDR3: CAKAAGSID) was found at expanded levels in the tracheas of one infection only individual and one single vaccination and infection individual. In the uninfected bird groups, 25 IgY public clones were present at expanded levels but absent or unexpanded in the infected groups (Figure 14). Interestingly, the identified public clones only show one or more expansions within the members of a single group, whilst generally not being detected outside of the particular immunisation regime where it was found. The only exception to this pattern is a clone (CDR3: CAKSAYGGYFGWGTYAGSID) which was found expanded in the spleen and bursa of a single vaccinated bird, whilst also being present at unexpanded levels in a double vaccinated bird. Additionally, clones identified in 6/11 of the double vaccinated group and 3/8 of the single vaccinated group were expanded across multiple individuals and tissues. By contrast, no clone that was identified in the naïve exhibited expansions across multiple individuals.

**Figure 13:**
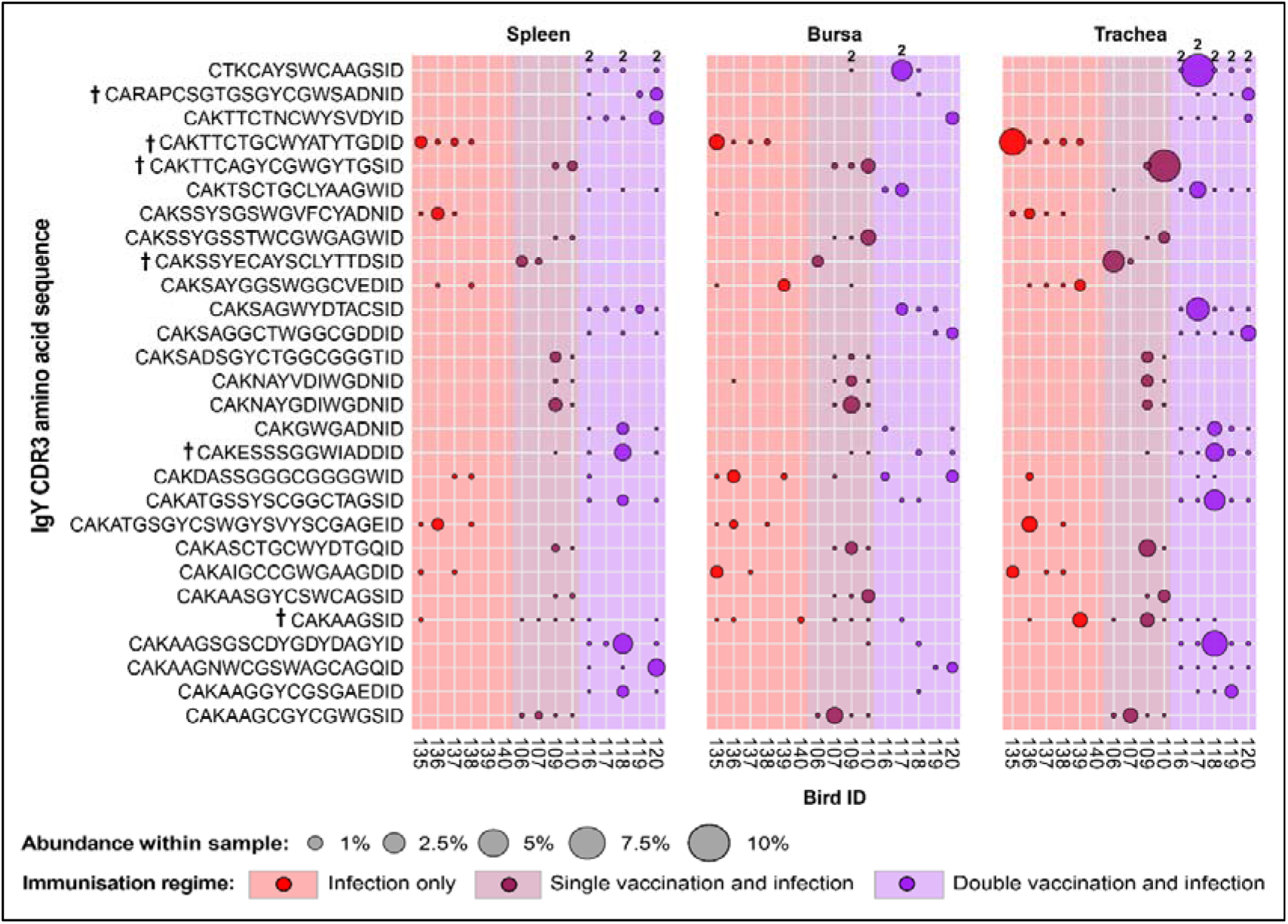
IgY clonal expansions in the restricted repertoire of infected birds. Circles indicate the presence of clones with a specific CDR3 amino acid sequence and are proportional to the abundance within each bird’s tissue clonal compartment. Overlapping circles display different clones based on nucleotide sequence which share the same CDR3 amino acid sequence. The plot shows only the clones which are expanded in the infected treatment groups (at or above 0.5% and at less than 0.5% in any of the uninfected). Clones which are expanded in multiple birds are identified with a dagger sign next to the CDR3 amino acid sequence. Background and circle colours indicate the immunisation regime identity: red – infection only; dark red – single vaccination and infection; purple – double vaccination and infection.

**Figure 14:**
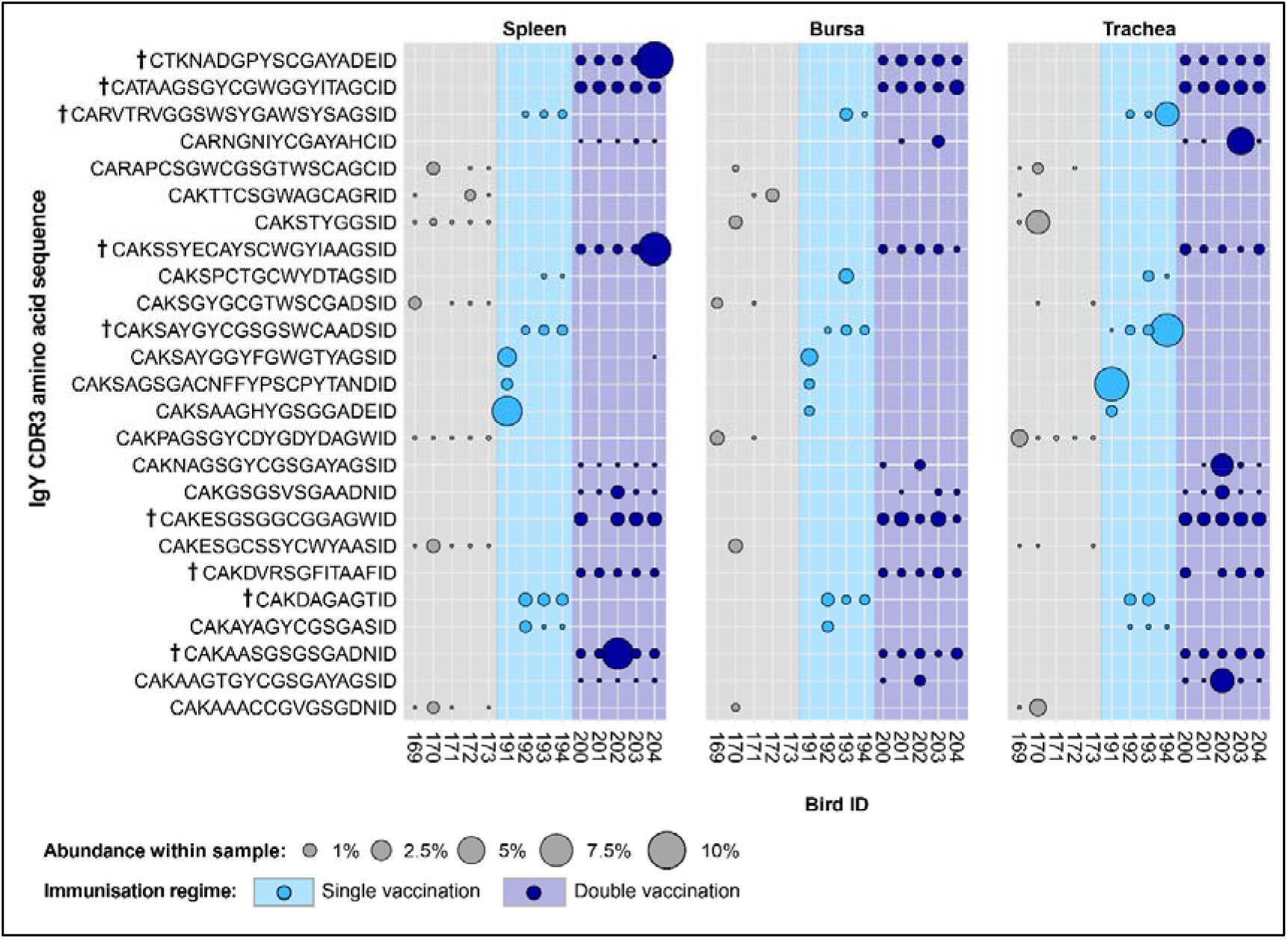
IgY clonal expansions in the restricted repertoire of uninfected birds. Circles indicate the presence of clones with a specific CDR3 amino acid sequence and are proportional to the abundance within each bird’s tissue clonal compartment. The plot shows only the clones which are expanded in the uninfected treatment groups (at or above 0.5% and at less than 0.5% in any of the infected). Clones which are expanded in multiple birds are identified with a dagger sign next to the CDR3 amino acid sequence. Background and circle colours indicate the immunisation regime identity: light blue - single vaccination; dark blue - double vaccination.

The majority of the 43 public IgY clones which were expanded in any of the vaccinated groups but below the expansion threshold or absent altogether from the unvaccinated groups (Figure 15), were either found previously as restricted to the infection immunisation regimes (19/43 - shown in red) or the uninfected treatment groups (20/43 shown in blue). The remaining 4 clones were expanded across both the infected and uninfected groups (shown in black). Only one of these clones (CDR3: CAKSAYGGSWGGFIEDID) was present across birds from all the vaccinated groups and exhibited expansions in individuals belonging to different immunisation regimes. The other 3 sequences were present either only in the single vaccinated groups (CDR3: CARAPCSTTWSCWYAAGSID) or only in the double vaccinated groups (CDR3s: CAKAARTAGYGVDDID and CAKAALTAGYGVDDID). Interestingly, the clones were present in the double vaccinated immunisation regimes only differ by a single amino acid and were found in all the birds of these groups. These clones were absent from some tissue samples and only expanded in the tissues of a double vaccinated bird and a double vaccinated and infected bird.

**Figure 15:**
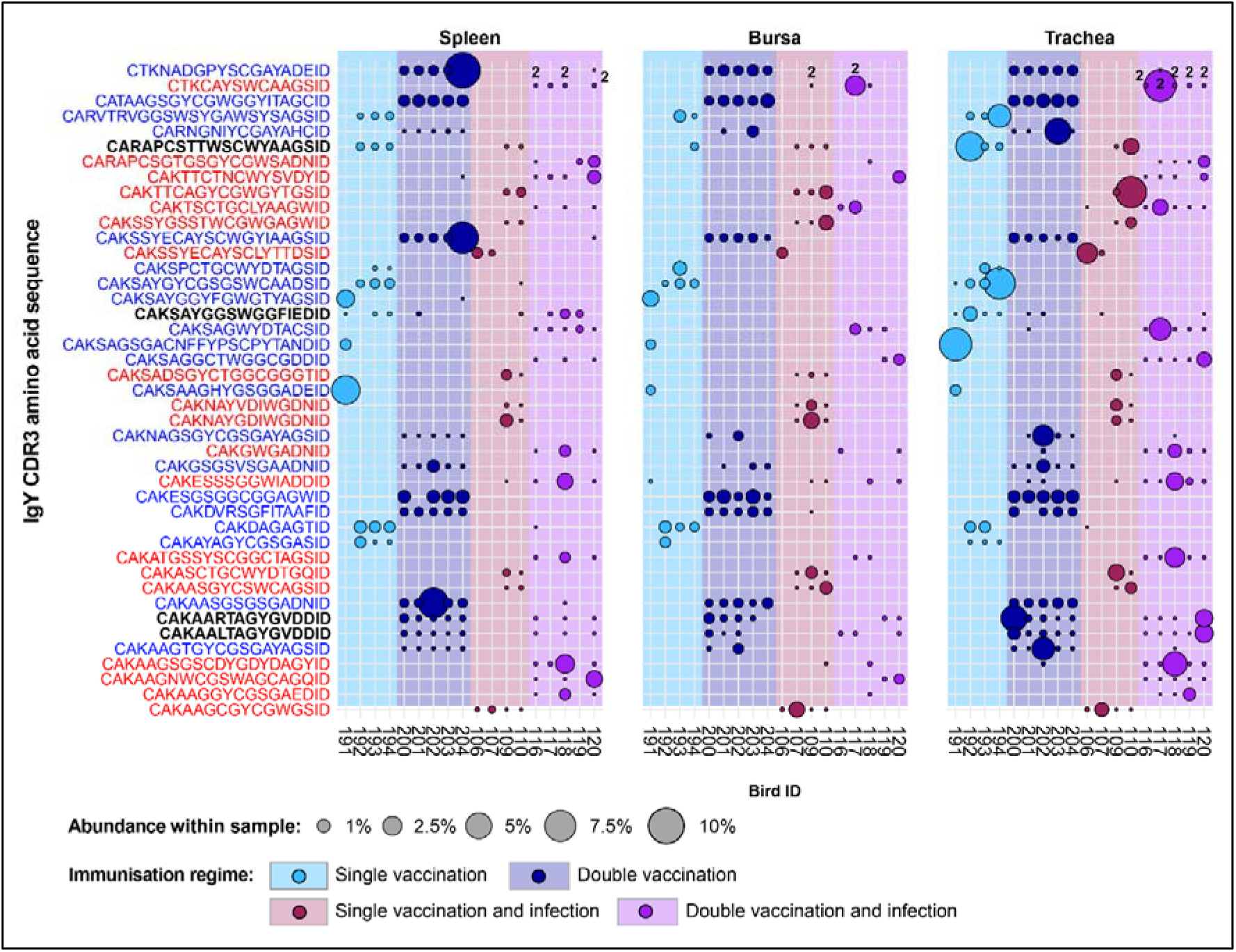
IgY clonal expansions in the restricted repertoire of vaccinated birds. Circles indicate the presence of clones with a specific CDR3 amino acid sequence and are proportional to the abundance within each bird’s tissue clonal compartment. Overlapping circles display different clones based on nucleotide sequence which share the same CDR3 amino acid sequence. The plot shows only the clones which are expanded in the vaccinated treatment groups (at or above 0.5% and at less than 0.5% in any of the naïve or infection only groups). CDR3 amino acid sequences which coincide with clones restricted to infected and uninfected birds are shown in red and blue, respectively. Background and circle colours indicate the immunisation regime identity: light blue - single vaccination; dark blue - double vaccination, dark red - single vaccination and infection; purple - double vaccination and infection.

## 4. Discussion

The antibody responses to AIVs are an important mechanism for viral clearance (27). However, the influences of viral infection or immunisation on the humoral immune system of birds are poorly understood, and the changes on the immune repertoire of H9N2-immunised chickens have not been previously explored. As the features of the avian immunoglobulin repertoire underpin the antigen specificity of the humoral response, information about the changes caused by antigenic exposure is key to both the understanding of the systemic antibody responses to infection and the protection offered by vaccination.

The patterns of expansion showed that the marked differences in the IgY and IgM repertoires of the birds are dependent both on immunisation regime and tissue type. Overall, the IgY repertoire exhibited higher clonal dominance than the IgM in the spleen and bursa, a pattern which is consistent with the antigen-specific expansion and class-switching, characteristic of IgY (28). Interestingly, the IgM repertoire in the trachea across all groups also exhibited higher dominance than in the bursa or spleen, showing levels that were slightly lower, yet comparable, to IgY. Moreover, the infected groups exhibited lower levels of IgY dominance in the trachea, which may indicate multiple clonal expansions in response to H9N2 infection in the upper respiratory tract. This pattern, although not as pronounced, was also observed in the IgM repertoires of the infected groups. Together, these differences suggest the resident IgM+ and IgY+ B cells are not as diverse in the trachea, but as class switching, expansion, and recruitment to the site of infection occur, the diversity of the IgY repertoire decreases with antigen-specific cells responding to infection. These observations are also supported by the diversity analyses, which revealed that at the level of dominant clones the infected groups are significantly more diverse both in IgM (double vaccination and infection) and IgY (infection only and single vaccination and infection).

The differences in diversity identified in the bursa and spleen may reflect a combination of tissue identity as well as the number and nature of the immunisations. In the bursa, the IgM repertoires displayed no differences in diversity between the groups. However, the IgY repertoires of the single vaccinated and the infection only groups exhibited significantly higher diversity levels than the naïve birds for clonal richness (D0), typical clones (D1), and dominant clones (D2). As the bursa is an organ not commonly associated with germinal centre development and class-switching to IgY, antigen-specific responses of resident IgY+ cells in the bursa constitute an interesting topic for future investigations. In the spleen, the differences between the groups in both IgM and IgY diversities were only statistically significant at the dominant clone (D2) level. The groups that received two or more immunisations, exhibited higher dominant clone diversity in either the IgY (double vaccination and double vaccination and infection) or IgM (single vaccination and infection) repertoires. Only the double vaccinated groups, irrespective of infection, exhibited higher dominant clone diversities in their IgM repertoires, whereas the single vaccination and infection group showed higher levels of IgY diversity which may reflect increased repertoire focussing with multiple immunisations. Lastly, with the trachea all immunised groups exhibited higher diversity levels for IgM and IgY when compared to the naïve birds, both in terms of clonal richness and typical clones. When tracheal dominant clones (D2) were considered, only infected groups showed higher levels of diversity in IgM (double vaccinated and infected) or IgY (infection only and single vaccinated and infected). This is most likely due to the local infection recruiting antigen specific B cells. Overall these data suggest that the nature of the immunisation(*s)* (vaccine and/or live infection) differentially affects the diversity of the BCR repertoire structure. Indeed, such differences in the BCR repertoire structure according to AIV vaccination have also been reported with humans (29,30).

A high degree of clonal sharing (publicness) between individuals was observed in both IgM and IgY repertoires, with the latter generally exhibiting higher proportions of public clones. The IgM and IgY public clones occupied significantly higher proportions of the repertoire than the private clones in the trachea, irrespective of immunisation regime. By contrast, the other tissues did not display such consistent differences between the public and private repertoires, with the exception of the IgM repertoires in the bursal samples, where all the groups, with the exception of the double vaccination and infection, had significantly higher proportions of private clones. This pattern is expected as the bursa is a site of BCR diversification in bird (5). However, no clear rule that can be attributable to the immunisation regimes seems to govern the public vs. private clonal compositions in the analysed bird tissues, but the data indicates that exposure to antigen by vaccination or infection influences these patterns. This might be expected if the naive repertoire was limited and exposure to antigen leads to expansion of similar/identical clones in different individuals. Public Influenza-specific BCR CDR3 amino acid sequences have been reported in humans, although many of these appear to arise from convergent selection rather than identical nucleotide sequences in different individuals (29,31). Furthermore, in humans, although the proportion of sequences attributed to these amino acids was relatively small, these clusters were larger than those attributed to private BCR (30).

The degree of clonal sharing among individuals revealed that the majority of the public compartment is comprised of rare publics, which were shared between two and up to 50% of the birds included in this analysis. Indeed, most of the previously identified differences in terms of (total) public clones were mirrored by the rare publics. Although the clonal compartments with higher degrees of clonal sharing, the common and the ubiquitous publics, occupied a much smaller proportion of the IgM and IgY repertoires, their proportional contribution indicates that a baseline of clonal sharing is present across individual birds. Furthermore, it is very likely that a higher sequencing depth would have resulted in clones which were deemed private by the current analysis to be found at low levels in multiple individuals and would thus be public. Together, the findings about clonal sharing have important implications, as they indicate that the diversity of immunoglobulin CDR3 specificities may be much more limited between multiple individuals than previously believed. This, in turn, might imply a constrain on an individual bird’s ability to respond to antigenic challenge, which has profound consequences in the defence against pathogens and efficient responses to vaccination. A possible explanation for the high degree of publicness observable in the IgM and IgY repertoires relates to the mechanism of immunoglobulin rearrangement during B cell development. While mammals use a RAG-dependent V(D)J rearrangement process, birds rely on gene conversion. Although, in theory, gene conversion can yield comparable or even higher magnitudes of CDR3 diversity, the genetic and physiological processes may in fact be constrained by or exhibit biases towards specific gene segments and/or towards specific CDR3 arrangements (31). This is an interesting hypothesis to explore, and future molecular studies are required to examine the gene conversion during immunoglobulin diversification in avian species.

Profound differences in the clonal expansion profiles restricted to specific treatments were evident between the IgM and IgY repertoires. Considerably fewer treatment- restricted expansions were present in the IgM repertoires as opposed to the IgY, a result which supports that antigen-stimulated IgM+ B cells undergo expansion and class-switching to IgY. Interestingly, for both IgM and IgY uninfected-restricted or infected- restricted expansions, the identified clones were generally only found in birds belonging to the same immunisation group. One explanation for this pattern is that environmental effects have impacted the bird groups differently and this has influenced their IgM and IgY repertoires. This may be true for the infected groups, as they were moved to isolators just prior to infection on day 21, and the environment, although expected to be similar to the initial conditions, was sufficiently different to exert this influence on their repertoires. However, the single vaccinated group and the double vaccinated group was housed in the same isolator, unlike the infection only group which was housed in a separate isolator. Similarly, all three of the uninfected groups were housed together throughout the experiment, including after the infection groups were moved to the isolators. Therefore, environmental differences alone could not account for the observed differences as groups that were housed together still exhibit expansions of the same clones which are not identified in other groups. Another explanation relates to experimental cross-contamination or day to day variation in process, but neither of these are likely since samples were batch processed for 5’ RACE PCRs and there were many group-specific features to the data. Since environmental factors, analytical variation, or cross-contamination were not able to account for the observed patterns, the interesting possibility remains that the specific combination of immunisation through infection and/or vaccination(s) may lead to convergent expansions of identical CDR3 between individuals.

The few IgY and IgM clones that were expanded across individuals belonging to multiple groups were restricted to infection, with none being shared across the uninfected groups. When the vaccination-restricted clones were considered, only one IgM clone and four IgY clones were found expanded in birds from the infected and uninfected groups. These identified CDR3 sequences restricted to the vaccinated groups and expanded in birds belonging to infected and uninfected groups may represent lineages of B cells that respond to H9N2 under both immunisation scenarios. These clones, stimulated through vaccination(*s*), may intrinsically possess a higher affinity for the virus, and could offer increased protection during infection. Furthermore, if a comparable number of restricted clonal expansions were to be detected in the IgL repertoires, screening for antigen specific (paired) light and heavy chain sequences (e.g. for the production of H9N2-specific monoclonal antibodies) might be narrowed down to a few antigen-specific light chain (IgL) and heavy chain (IgH) CDR3 sequences based solely on repertoire data. Although this was outside the scope of the current research, such endeavours could substantially increase the throughput in generating antigen-specific antibodies not just for avian influenza viruses, but for other infectious agents as well.

In conclusion, our analyses of the IgM and IgY repertoires of chickens subjected to different H9N2 immunisation regimes revealed important findings which constitute a solid foundation for future research aimed at increasing our understanding of the avian humoral responses to avian influenza, and the adaptive immune system more broadly. The findings presented herein strongly suggest that not only does the nature of the immunisation (i.e. vaccination or infection) influence the immunoglobulin repertoires of the individuals, but also the number of immunisations received and the particular combination of immunisations to which the birds were subjected to. Unravelling the complexities of the avian immunoglobulin repertoires thus serves as a fruitful area for future research, whilst also having the potential to inform on practices such as vaccination, which remains paramount for efficient infectious control at a global level.

## Acknowledgements.

The authors would like to acknowledge the Pirbright Institute Flow Cytometry unit and support through the Core capability grant (BBS/E/I/00007039). The authors are also very grateful to Oenone Bodman-Harris for proofreading and providing useful suggestions for the final version of the manuscript.

## Supplementary Figures and Legends

**Supplementary Figure 1:**
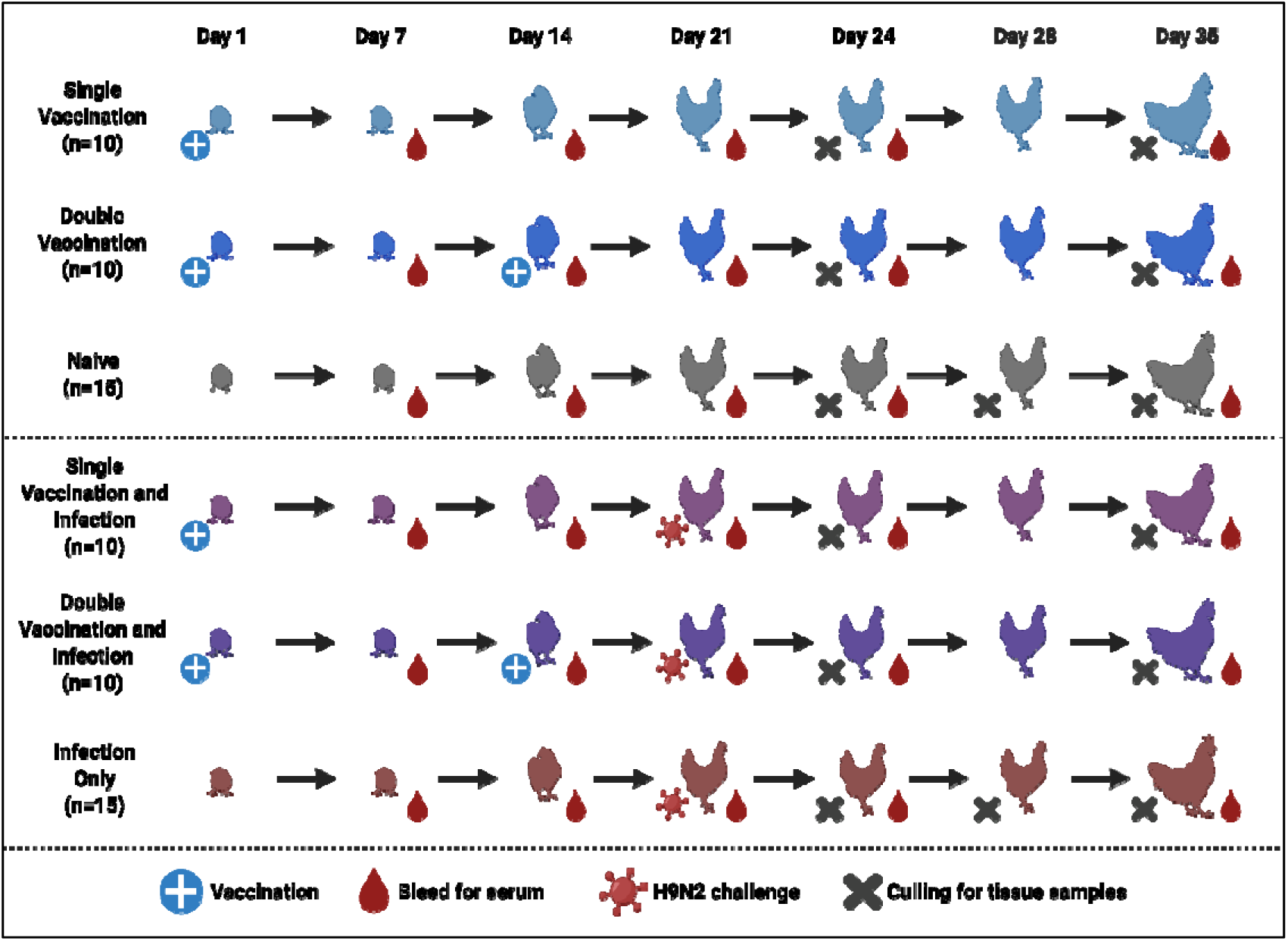
Design of the H9N2 vaccination and infection experiment. Birds were split into 6 groups which received either an inactivated H9N2 vaccine at day 1, or both at days 1 and 14, or no vaccination. At day 21, half of the birds (belonging to all vaccination regime treatments) were infected with H9N2 avian influenza virus. Birds were then culled at days 24 (n=5 from all groups), day 28 (n=5 from the unvaccinated treatments), and at day 35 (n=5 from all groups). Blood and tissue samples were harvested and processed for subsequent analyses. Buccal and cloacal swab samples were collected from the infected birds with one pre-infection sampling and 10 other daily swabs after infection.

**Supplementary Figure 2:**
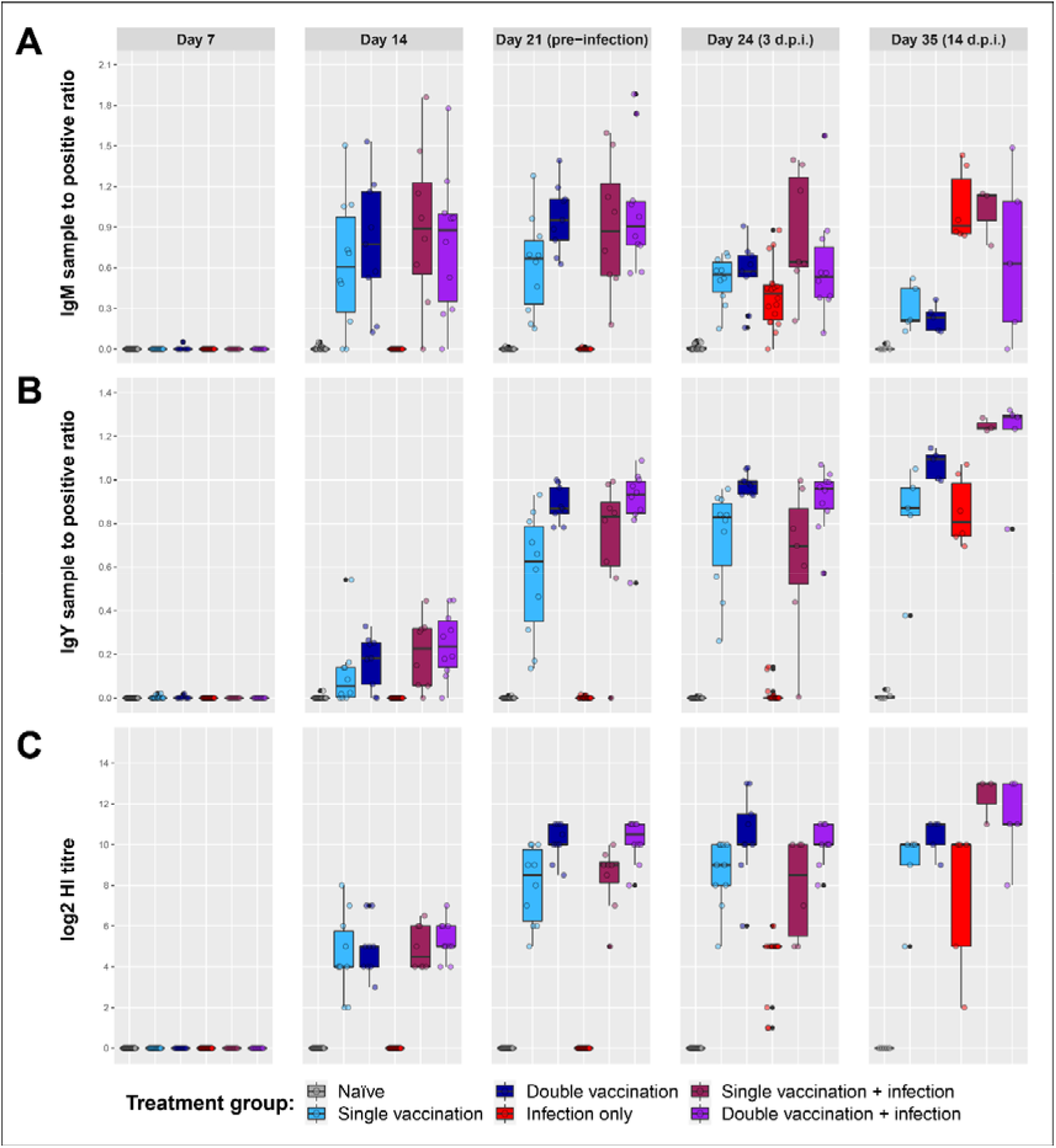
H9N2-specific antibody levels and haemagglutination inhibition (HI) potential of sera in chickens following vaccination and infectious challenge. **(A)** IgM ELISA sample-to-positive ratios. **(B)** IgY ELISA sample-to-positive ratios. **(C)** HI titres of serum samples.

**Supplementary Figure 3:**
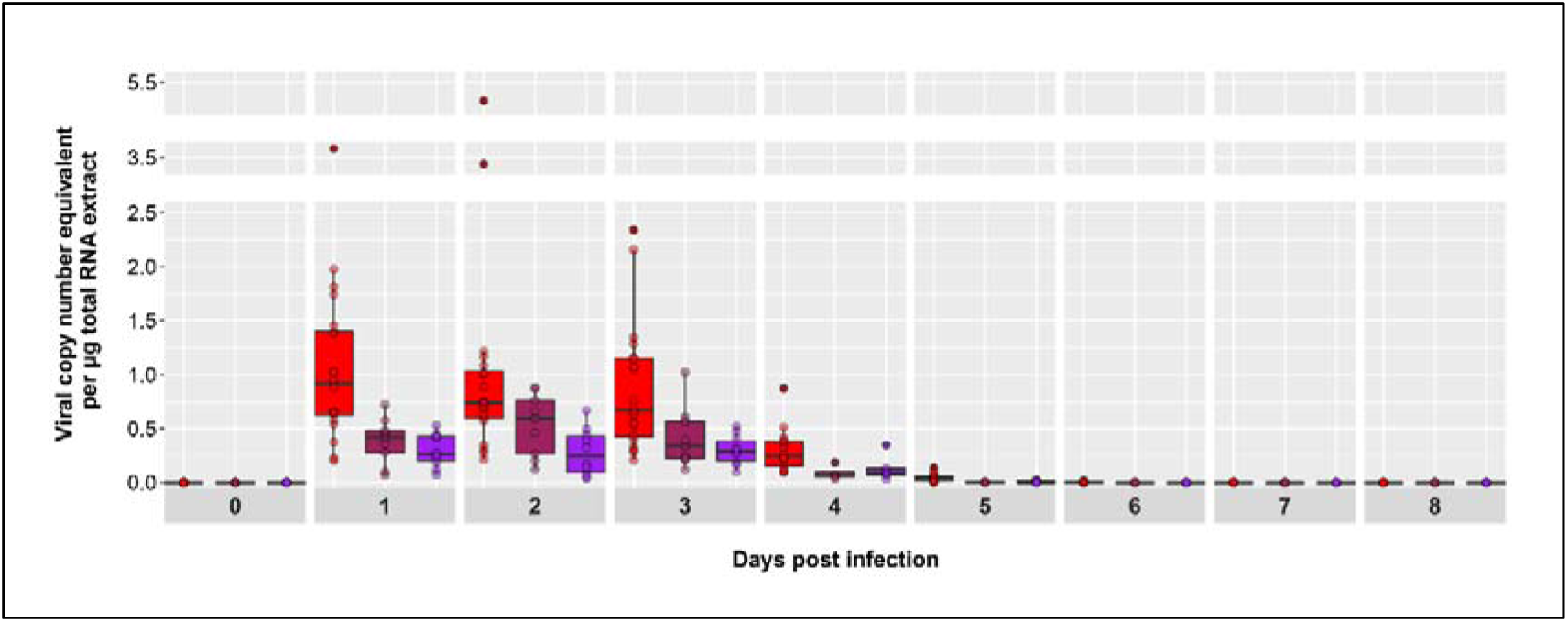
H9N2 matrix gene qRT-PCR results of buccal swab samples. Viral copy number equivalent per μg total RNA extract was calculated for the infected bird groups from day 0 (pre-infection) until day 8 post-infection.

**Supplementary Figure 4:**
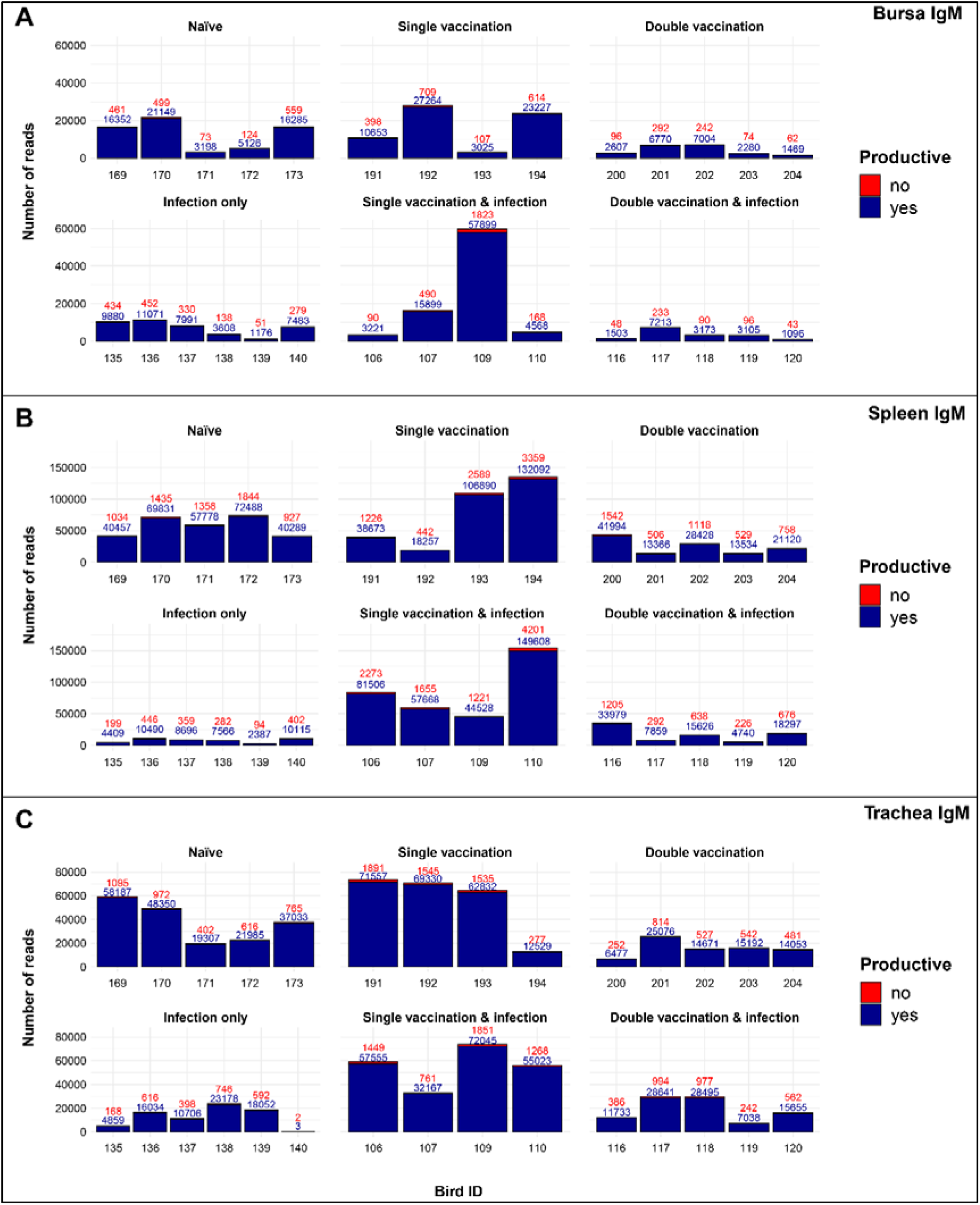
Total number of IgM sequence reads identified in tissues of chickens that were subjected to different immunisation regimes. (A) Splenic samples, (B) bursal samples, (C) tracheal samples. Bird numbers displayed on the x axis and individuals are grouped based on the corresponding immunisation status which is illustrated above each panel. Productive and unproductive reads are shown in blue and red, respectively.

**Supplementary Figure 5:**
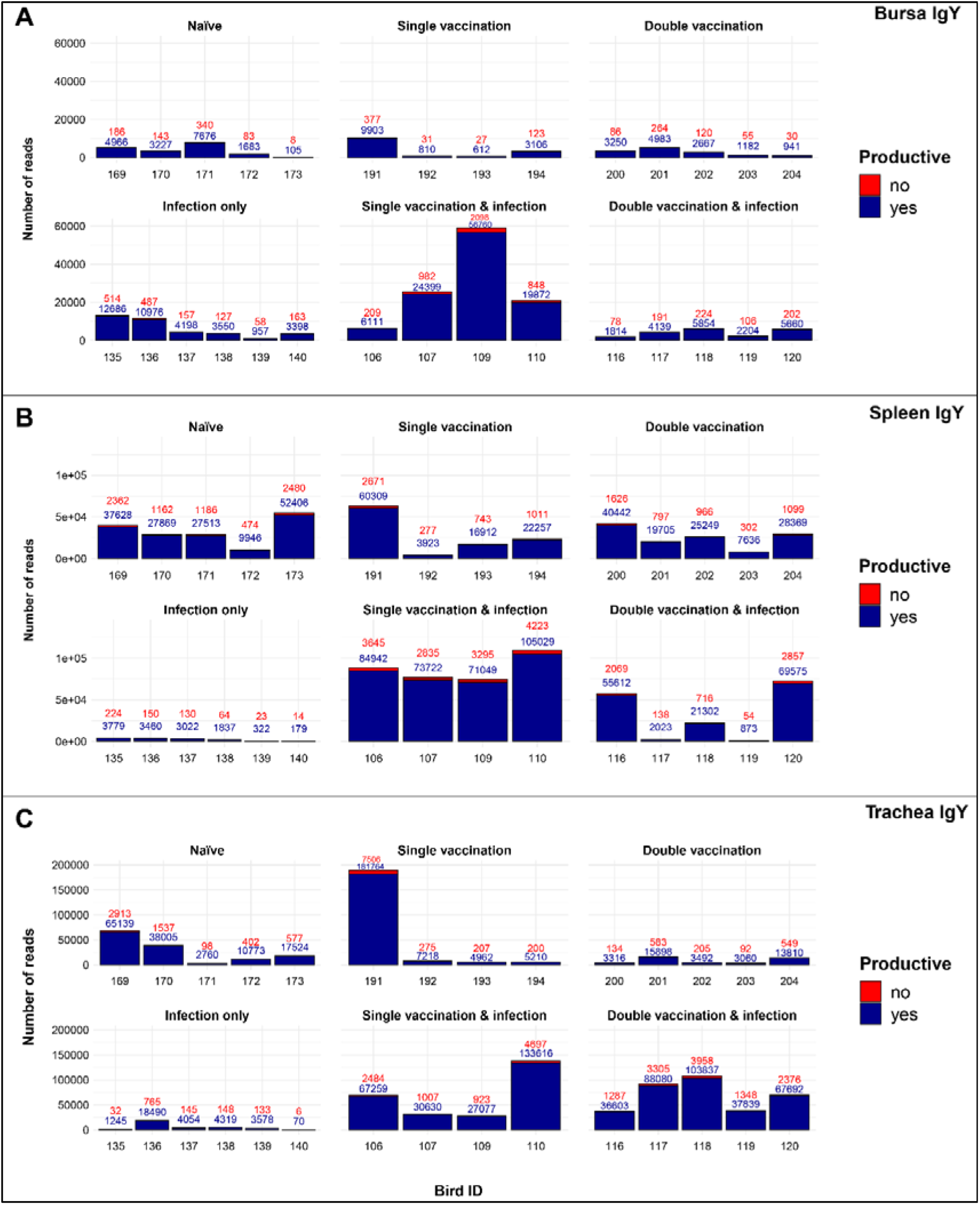
Total number of IgY sequence reads identified in tissues of chickens that were subjected to different immunisation regimes. (A) Splenic samples, (B) bursal samples, (C) tracheal samples. Bird numbers displayed on the x axis and individuals are grouped based on the corresponding immunisation status which is illustrated above each panel. Productive and unproductive reads are shown in blue and red, respectively.

